# Microglia undergoes chemokine receptor, CX3CR1-mediated internalization of extracellular Tau

**DOI:** 10.1101/2024.12.03.626734

**Authors:** Hariharakrishnan Chidambaram, Smita Eknath Desale, Kapali Suri, Arul Murugan N, Subashchandrabose Chinnathambi

**Affiliations:** Neurobiology Group, Division of Biochemical Sciences, CSIR-National Chemical Laboratory, Dr. Homi Bhabha Road, 411008 Pune, India; Academy of Scientific and Innovative Research (AcSIR), Ghaziabad, 201002, India; Department of Neurochemistry, National Institute of Mental Health and Neuro Sciences (NIMHANS), Institute of National Importance, Hosur Road, Bangalore, 560029, Karnataka, India; Department of Computational Biology, Indraprastha Institute of Information Technology, New Delhi 110020, India; Center for Excellence in Healthcare, Indraprastha Institute of Information Technology, New Delhi 110020, India

**Keywords:** Extracellualr Tau, Chemokine receptor CX3CR1, Microglia, Tau internalization, cell migration

## Abstract

In Alzheimer’s disease, the microtubule-associated protein, Tau misfolds to form aggregates and filaments in the intra- and extracellular region of neuronal cells. CX3CR1 receptor is involved in monomeric Tau internalization by directly binding to the receptor. In our study, we are interested in understanding the interaction of different pathological structures of Tau with microglial chemokine receptor, CX3CR1 and its role in CX3CR1-mediated Tau internalization by immunofluorescence assay. Co-immunoprecipitation further confirmed the receptor interaction with Tau oligomers and even aggregated filaments. We next determined the role of Tau as a chemoattractant in regulating the migration of microglia by wound healing assay. Further, immunolocalization studies of CX3CR1 with β-arrestin proved the desensitization of CX3CR1 receptor upon Tau binding. Upon ligand interaction, GPCRs undergoes desensitization and encytosed to the cytosol. We have quantified the CX3CR1 internalization by measuring the membrane to cytosol intensity ratio of CX3CR1 receptor upon extracellular Tau exposure. In addition to experimental assays, a comprehensive analysis of the interaction between the CX3CR1 receptor, the Tau protein, and the AZD8797 ligand has been derived from the combined molecular docking, molecular dynamics and free energy calculations using molecular mechanics-generalized Born approach. In particular, two different molecular dynamics simulations were carried out for CX3CR1-Tau protein and AZD8797-bound CX3CR1-Tau protein interaction. The molecular modeling study establishes that AZD8797 binds to CX3CR1 receptor and modulates the Tau interaction with the receptor.

## Introduction

Alzheimer’s disease is a neurodegenerative disorder that progressively leads to cognitive and behavioral impairment [1–3]. The intracellular accumulation of misfolded, self-aggregated Tau fibrils and extracellular senile plaques from Amyloid-β peptides are the pathological hallmarks of Alzheimer’s disease [2]. Tau is a microtubule associated protein expressed in the axon of neuronal cells involved in the microtubule stabilization of the central nervous system [4,3,5]. Tau is soluble, cytoplasmic and natively unfolded protein that binds at the interface of α-β tubulin heterodimers of axonal microtubule [4]. Human full-length Tau consists of 441 amino acid residues with two N-terminal inserts, N1, N2, a proline-rich domain and four repeat regions, R1, R2, R3 and R4 [6]. The repeat region of Tau is majorly involved in microtubule binding with two flanking N-and C-terminals. In neurodegenerative diseases such as Alzheimer’s disease, Tau undergoes several post-translational modifications mainly phosphorylation and detaches from microtubules to form oligomers and aggregated fibrils in the cytosolic region [7,8]. The misfolded and accumulated Tau species escape and spread through leaky substances from dying neurons, exosomes, *etc* and implants seeding effect in the neighbouring healthy neuronal cells [9]. Microglia are the brain macrophage cells that actively surveys the brain microenvironment and phagocytose foreign materials such as pathogens, cell debris, apoptotic cells, misfolded protein aggregates, *etc* [10]. Microglia sense these extracellular particles through a wide variety of cell surface receptors such as ion channels, G-protein coupled receptors, transmembrane receptors, *etc*. In Alzheimer’s disease, microglia plays a dual role of both protecting and destroying the neuronal cells based on the stage of disease progression. In earlier stages of AD, microglia is involved in the clearance of extracellular Tau fibrils and Amyloid-β plaques [11–15]. Whereas, at later stage of disease progression, microglia gets constantly activated, and phagocytose neuronal cells that ultimately leads to neuroinflammation and neuronal damage [16,17]. Microglia internalizes extracellular Tau by various mechanisms such as phagocytosis, micropinocytosis, clathrin and dynamin-dependent endocytosis that either accumulates in the cytosol or released via exosomes or degraded by lysosomal fusion [18–21]. Several membrane receptors are involved direct binding to extracellular Tau species such as purinergic P2Y12 receptor, chemokine CX3CR1 receptor, LRP1 receptor, M1 and M3 muscarinic acetylcholine receptors and promotes receptor-mediated internalization of extracellular Tau [12,22–25]. These receptors are majorly studied for their cross-talk with misfolded Tau protein that may lead to alternative downstream-signalling, up-regulate/ down-regulate receptor expression, change in receptor localization, *etc*. Chemokine receptor, CX3CR1 is a class A family of G-protein coupled receptors. CX3CR1 receptor is majorly involved in cell migration of white blood cells such as lymphocytes and monocytes [26]. In the brain, CX3CR1 receptor is abundantly expressed by microglia followed by minimal expression in synaptic terminals of neuronal cells [27,28]. In microglia, CX3CR1 receptor has its key role in cell adhesion and migration [29,30]. The ligand of this receptor, fractalkine (CX3CL1) is expressed by neurons and astrocytes and distributed at the cell membrane with its mucin-like stalk and N-terminal receptor-binding domain extracellularly for receptor binding [31,32]. Microglia maintains a resting state upon binding to the membrane-bound fractalkine of neuronal cells. Upon cleavage by metalloproteases, the soluble form of fractalkine (Mucin-like stalk and the chemokine domain) detaches from the membrane [33,34]. This soluble fractalkine acts as a chemoattractant that promotes microglial migration towards the site of neuronal inflammation [35]. CX3CR1 deficiency exacerbated the disease condition with increased Tau phosphorylation showing the significance of this receptor in regulating neuroinflammation. This activity was dependent on functional toll-like receptor 4 (TLR4) and Interleukin-1 receptor (IL-1) with an increased activity p38 MAPK [36]. In 2017, Bolos *et al*. first studied the role of microglial CX3CR1 receptor in Tau internalization [12]. In AD conditions, Tau oligomers are considered as the most toxic species and rapidly promotes further Tau oligomerization and aggregation [37]. In this study, we are interested in determining the role of chemokine receptor, CX3CR1 interaction with different pathological structures of Tau such as oligomers and aggregates by direct binding. CX3CR1-mediated Tau internalization is performed in the presence and absence of CX3CR1 antagonist, AZD8797. We have also studied the role of Tau species as a chemoattractant in promoting the migratory state of microglia. Upon ligand binding, GPCRs undergo desensitization by receptor phosphorylation and β-arrestin recruitment. CX3CR1 receptor undergoes receptor desensitization upon activation which is mediated by GRK-2 phosphorylation and β-arrestin recruitment to the receptor, ultimately endocytosed for either degradation or receptor recycling [38]. In the later part, we have studied the Tau-mediated CX3CR1 desensitization by immunolocalization studies with β-arrestin and proved the internalization of CX3CR1 by intensity profiling of microglial membrane.

## Experimental procedures

### Chemicals and reagents

Himedia - Luria-Bertani broth; MP Biomedicals - MgCl_2_, KCl, NaCl, Na_2_HPO_4_, KH_2_PO_4_, ethylene glycol-bis (baminoethylether)- N,N,N′ ,N′ -tetraacetic acid (EGTA), phenylmethylsulfonylfluoride (PMSF), ammonium acetate, sodium azide, ammonium persulfate (APS), methanol, Dimethyl Sulfoxide Solvent (DMSO) and ampicillin Calbiochem - Isopropyl β-d-1- thiogalactopyranoside (IPTG) and dithiothreitol (DTT). Invitrogen – Pierce co-immunoprecipitation Kit (Thermo 23600), Alexa 647-C2 maleimide, Paraformaldehyde, Pierce™ Tris(2- carboxyethyl)phosphine hydrochloride (TCEP-HCl), Pierce™ RIPA buffer, Tris base, acrylamide, and N,N,N′,N′ -Tetramethyl ethylenediamine (TEMED). Merck - Immobilon PVDF membrane. Biorad - Precision Plus Protein Standards (Bio-Rad, cat no. 161–0374), Glycine and Protein Assay Dye Reagent. Sigma - CuSO4, bicinchoninic acid (BCA), bovine serum albumin (BSA), Thioflavin S (ThS), MES, BES, TritonX-100, glutaraldehyde, and Sodium dodecyl sulphate (SDS). Roche cOmplete™ - Protease inhibitor cocktail.

### Cell culture requirements

Invitrogen - RPMI 1640 media, PBS buffer (1X), Fetal Bovine Serum (FBS), Horse serum, Penicillin-streptomycin, and trypsin-EDTA. **Antibodies:** CX3CR1 rabbit monoclonal antibody (Thermo-702321), CX3CR1 mouse monoclonal antibody (Santa cruz sc-377227), β-arrestin-1 mouse monoclonal antibody (Thermo 39-5000), Total pan-Tau antibody K9JA (Dako, A0024), β-Actin loading control monoclonal antibody (BA3R) (Thermo, MA5–15739), Phalloidin-Alexa 488 for F-actin (Thermo, A12379), Goat anti-rabbit IgG (H+L) Cross-adsorbed secondary antibody HRP (Invitrogen, cat no. A16110), Antimouse secondary antibody conjugated with Alexa flour-488 (Invitrogen, cat no A-11001), Goat anti-rabbit IgG (H+L) Cross-adsorbed secondary antibody with Alexa Fluor 555 (Invitrogen, cat no. A-21428), and DAPI (Invitrogen, cat no. D1306). R&D systems - Recombinant fractalkine/CX3CL1 (Catalogue no. 458-MF/CF). MedChemExpress – Human CX3CR1 antagonist, AZD8797.

### Tau protein expression and purification

Human-Tau protein is recombinantly expressed and purified as per the protocol mention in Chidambaram *et al*. 2022 [39]. In brief, human full-length Tau is recombinantly expressed in BL21* strain of *E.coli* cells by inducing with 0.5mM isopropylthio-β-galactoside (IPTG). Cells are harvested after 4 hours incubation and lysed in cell disruption system using cell lysis buffer containing 20 mM MES, 0.2 mM MgCl2, 1 mM EGTA, 1 mM PMSF, 5 mM DTT, and PIC (0.01%). The lysate is then added with excess NaCl and boiled at 90°C for 15 minutes. The supernatant is collected by centrifuging the lysate at 40000 rpm for 45 minutes at 4°C and excess NaCl is removed by dialysis for overnight at 4°C. Tau protein is further purified by cation exchange chromatography, followed by gel-filtration chromatography. The concentration of the protein is estimated and stored at -80°C for further studies.

### Tau oligomer and aggregate setup

Tau oligomers are prepared in PBS buffer pH.7.4 containing 2 mM DTT, 0.01% Sodium azide, 25 mM NaCl, and 1X PIC. Heparin 17,500 Da is added to Tau in the ratio of 1:4 and incubated at room temperature for 12 hours. Oligomers are cross-linked with 0.01% glutaraldehyde and purified by size-exclusion chromatography. Tau aggregates are prepared in 20mM BES buffer pH.7.4 containing 1 mM DTT, 0.01% Sodium azide, 25 mM NaCl, and 1X PIC. Heparin 17,500 Da is added to Tau in the ratio of 1:4 and incubated at 37 °C for 72 hours. The concentration of the protein is estimated and stored at -80°C for further studies. The quality of Tau oligomers and aggregates are determined by SDS-gel electrophoresis, ThS fluorescence assay, ANS assay and transmission electron microscopy (TEM) analysis [21].

### Tau species labelling using Alexa-fluor 647 C_2_ maleimide

100 µm monomer, oligomer, and aggregate are labelled with Alexa-fluor 647 C2 maleimide in the ratio of 2:1 (C2 maleimide: Tau) and incubated at 4°C shaking (600 rpm) overnight. Buffer exchange was performed with PBS buffer pH.7.4 using 10-kDa centrifugal filters to remove excess dye from the solution. Monomeric Tau is added with 2µM Tris (2-carboxyethyl) phosphine (TCEP) and incubated for 10 minutes at shaking conditions (600 rpm) before adding the dye to maintain the cysteine residues of Tau at reduced conditions. The concentration of the protein is estimated and stored at -80°C for further studies. The quality of labelled Tau monomers, oligomers and aggregates are determined by SDS-gel electrophoresis, ThS fluorescence assay, ANS assay and transmission electron microscopy (TEM) analysis [22].

### Cell culture

N9 microglia are revived in RPMI media containing 10% fetal bovine serum and incubated with 5% CO2 supply at 37°C. Cells are passaged at 80% confluency and seeded in multi-well plates for the experiment. The N9 microglial cell line no. is CVCL_0452.

### Immunostaining of N9 microglia

N9 microglial cells are harvested and counted from the culture dish. 20,000 cells per well are seeded in a 24 well sterile dish containing coverslips and incubated for 24 hours at 37° incubator with a 5% CO_2_. RPMI treatment media (Reduced serum media) containing 1% FBS is added to the cells. Alexa-fluor labelled Tau species (Monomer, oligomer and aggregate) are added at desired concentration (1µM) and incubated for different time intervals 5, 30, 60 and 180 minutes. Internalized Tau intensity, internalization and localization of CX3CR1 receptor, and β-arrestin were studied from the immunofluorescence microscopy analysis. In inhibitor-based studies of CX3CR1 receptor, AZD8797 (CX3CR1 antagonist) is provided as a pre-treatment to the microglial cells at 100 nM concentration. Fractalkine at 250 ng/mL is provided to microglial cells for receptor activation. Following the treatment the cells are fixed using 4% paraformaldehyde for 20 minutes at room temperature. The coverslips are rinsed once with PBS buffer and permeabilized with PBS buffer containing 0.2% triton-X (PBS-t) for 20 minutes at rocking conditions. The coverslips are then blocked using PBS-t buffer containing 5% horse serum for 1 hour at room temperature. Atlast, the coverslips are incubated desired with primary antibodies at 4°C overnight with humid conditions, followed by secondary incubation at room temperature for 1 hour. 3 times of PBS-t wash for 10 minutes is provided after the antibody incubation at room temperature. **Antibody dilutions.** CX3CR1 Recombinant Monoclonal Antibody (702321) - 1:200; β-arrestin-1 mouse monoclonal antibody (39-5000) – 1:200; Goat anti-mouse secondary antibody conjugated with Alexa Fluor 555 – 1:500 and Goat anti-rabbit IgG (H+L) Cross-adsorbed secondary antibody with Alexa Fluor 488 – 1:500. The coverslips are incubated with 300 nM DAPI for 10 minutes to stain the nucleus. The coverslips are mounted on clean glass slides using Prolong Diamond Antifade mountant and cured for 24 hours at room temperature.

### Fluorescence microscopy

The coverslips are observed under Zeiss Axio observer Z1/ 7 Apotome 2.0 wide-field fluorescence microscope with Plan Apochromat 63x/ 1.40 Oil DIC M27 objective lens. Axiocam 503 is used for image-capturing with fluorescence intensities from different targets of the cell. The intensity and exposure time are set constant as per the experiment and the images are acquired as .czi files. Z-stack images are acquired at desired fields and all the images are uniformly processed by adjusting the black and white balance of the fluorescence intensity in Zen 2.3 software. The experiments were repeated for three independent repeats (n=3). **Colocalization.** The extent of colocalization between Tau, CX3CR1, and β-arrestin are calculated from Pearson’s colocalization coefficient. The colocalization between two different units (Pearson’s co-efficient) is calculated using Coloc2 plugin of ImageJ 1.51s software. **Intensity quantification.** The internalized Tau intensity is calculated from the immunofluorescence images by contour tool of Zen 2.3 software that provides intensity mean values of different fluorophores from a desired region of interest. **Intensity profiling.** The surface to cytosolic intensity ratio of CX3CR1 receptor is calculated by the profile tool from graphics tab of Zen 2.3 software. A line of ∼6µm is drawn from the cortical membrane towards the cytosol. The profile graph shows the intensity values of all fluorophores that the line crosses through from the field. The intensity profile is also used to show the colocalization between Tau and CX3CR1 receptor in individual treatment groups.

### Co-immunoprecipitation

N9 cells are passaged and seeded in 3-5 100 mm culture dish for co-immunoprecipitation assay. The cells are incubated at 37° with 5% CO2 supply until they reach the confluency. The co-immunoprecipitation assay was performed as per manufacturer’s protocol with few modifications. First, the cells were harvested with CO-IP lysis buffer upto 1 ml per 100 mm culture dish and incubated for 30 minutes with pipetting at regular intervals. Upon lysis, the lysate is centrifuged at 14,500 rpm for 10 minutes to remove any cell debris, nucleus, *etc*. Pan-Tau K9JA primary antibody was coupled with the amino-linked resins by a reductive amination reaction as per manufacturer’s protocol. Appropriate controls for non-specific binding and isotype IgG were maintained. Tau species (1 µM and 5 µM) is then added to the antibody-coupled resin columns and incubated at 4 °C rotor for 2.30 hours. Excess Tau species is removed by spinning the column at 3000 rpm for 60 seconds. An equal volume of lysate was added to every Tau containing (1 µM and 5 µM) K9JA-coupled resin columns and one group without Tau (with K9JA-coupled resin) as a negative control. The mixture is then incubated by constant shaking for 15 minutes (RT). Multiple washes with wash buffer, followed by conditioned buffer was performed in-order to remove the non-specific protein binding as per the protocol. Proteins that interacted with Tau were eluted using the elution buffer according to the manufacturer’s protocol. The eluted proteins are analysed by SDS-PAGE and western blot by specific protein and K9JA antibody (CX3CR1-1:200 and K9JA-1:8000 dilution).

### Western blot

CX3CR1 expression and the total Tau internalization (Monomer, oligomer and aggregate) have been confirmed by western blot analysis. N9 microglial cells are counted and 3 lakh cells per well were seeded in sterile multi well dish and incubated for 24 hours at 37° with 5% CO_2_. The cells are replaced with RPMI treatment media (reduced serum media) with 1% FBS. Microglia cells are then exposed to different Tau species at 1µM concentration for different time intervals. For CX3CR1 inhibition and activation, 100nM AZD8797 and 250ng/mL fractalkine are added as pre-treatment respectively. Cells are harvested by trypsinization and pelleted down by centrifuging at 800 rpm for 5 minutes. The cells were lysed as per the protocol mentioned by Chidambaram et al 2022. In brief, the cells were lysed with RIPA lysis buffer containing 1% Triton-X and 1X PIC. The lysate is centrifuged to remove the nucleus and other debris at 14,500 rpm for 20 minutes at 4°C. The supernatant protein concentration is estimated by Bradford assay and an equal load of lysate was loaded in 10% SDS-gel electrophoresis. The protein is transferred to methanol activated PVDF membrane using Amersham semi-dry transfer unit. The current is set constant at 200mA and the transfer time is adjusted according to the molecular weight of the protein of interest. After transfer, the membrane is blocked using 10% skimmed milk solution in PBS buffer containing 0.1% Tween-20 (PBS-t). The membrane is incubated in primary antibody overnight at 4°C/ 1 hour at 37°C, followed by secondary incubation for 1 hour at room temperature, shaking conditions. (Dilutions: K9JA antibody – 1:8000; CX3CR1 mouse monoclonal antibody – 1:200; β-actin rabbit monoclonal antibody – 1:2500; Goat anti-rabbit IgG (H+L) Cross-adsorbed secondary antibody HRP – 1:10,000; Goat anti-mouse IgG (H+L) Cross-adsorbed secondary antibody HRP – 1:1500). 3 times of PBS-t wash for 10 minutes is provided after the antibody incubation at room temperature. The blot is developed with SuperSignal™ West Atto Ultimate Sensitivity Substrate in chemiluminescence mode of Amersham Imager 600. The intensity of the desired protein band is quantified using Image Lab software version 6.0.1 and normalized using β-actin loading control. The experiments were repeated for three independent repeats (n=3). The fold change in the expression of CX3CR1 and the internalized Tau levels were confirmed by western blot.

### Wound-healing assay

N9 microglia cells were seeded at a density of 5*10^5 cell/well and the scratch was made with 200 µl pipette tip to the confluent monolayer of the cell. After scratch the cells were given two washes of 1X PBS to remove unattached cells from the monolayer. Followed by the washing, extracellular treatment of Tau monomer, oligomer, aggregates (1 µM), and AZD (100 nM) was given to cells in combination; positive control of Fractalkine (250 ng/ml) was kept to compare the assay. The migration of cells into the wound from both the sides was observed at 0, 6, 12, 24 hours for all the treatment condition. The migration of cells was observed at 10X magnification in Zeiss phase-contrast microscope.

### Statistical analysis

All experiments are performed in three biological replicates, and triplicate measurements are taken for each individual experiment. The statistical significance is determined by one-way ANOVA using GraphPad Prism 8.0.1. For multiple groups, the significance is calculated by Tukey-Kramer’s post-hoc analysis at 5% level of significance. The p values are then calculated by comparing with control group and the level of significance is mentioned within the graph (*** p<0.001, ** p<0.01, * p<0.05, ns p≥0.05). The results are considered significant if the mean difference between treatment groups is greater than calculated Tukey’s criterion (X-X’>T).

### Preparation and Docking of the protein structure with the CX3CR1-ligand (AZD8797)

The 3D structure of the CX3CR1 receptor was built using Modeller software [40]. The sequence was retrieved using accession number AAY21515.1 from the NCBI database (https://www.ncbi.nlm.nih.gov/). Further, blastp was used to identify the protein structure with the highest degree of similarity with the sequence of the receptor. The PDB structure with PDB ID 8IC0 was chosen as the template as it had the highest percentage of identity (100%) and the lowest E value (0). After utilizing Modeller to perform homology modeling, the model with the lowest DOPE (Discrete Optimized Protein Energy) score was chosen for docking. The repeat domain of the Tau model created by Sonawane *et al*., 2019 [41] in earlier works was used for further interaction investigation. The Electrostatic Potential analysis was carried out for both CX3CR1 receptor and Tau using Pymol to analyze the distribution of charges to get insight into how both receptor and Tau interact with each other depending on their electrostatic potential. Using Uniprot, the CX3CR1 receptors’ extracellular domains were determined. In the extracellular region, active residues were identified using Castp3.0 [42]. Additionally, utilizing the Castp-identified active sites, the HADDOCK [43,44] program was employed to carry out protein-protein docking for receptor with the Tau protein. The CX3CR1 antagonist, AZD8797 was docked with this above complex using Autodock vina [45] to investigate how the molecule modulates the interaction between the receptor-Tau complex. The docked complex in **Figure. 5B** was analyzed using Pymol and then Amber MD simulation [46] was performed for the binary and ternary complex to explore the structure, stability, binding energetics and conformational dynamics of the CX3CR1, Tau binary complex and CX3CR1, Tau protein, AZD8797 ternary complex.

### Molecular Dynamic Simulation

The process of preparing a molecular system for molecular dynamics simulation using AmberTools involves several sequential steps, encompassing parameterization, topology generation, and solvation. The ligand preparation was performed using AmberTools, by utilizing the GAFF force field [47] which is suitable to describe the organic molecules. The charges for ligand were generated using B3LYP/6-31G* level of theory by employing Gaussian09 software [48]. In general, the force-field development for ligand involves conversion of the mol2 or PDB file to create a prepi file and frcmod file using antechamber and Parmchk2 modules. We have used FF99SB force field [49] for the receptor and Tau protein. The water solvent was described using TIP3P force-field. The receptor-Tau protein complex and receptor, Tau protein with AZD8797 ternary complex were embedded in the lipid membrane for which the Charmm-GUI [50] was used. The binary and ternary systems were neutralized with counter ions and a sufficient number of water solvents were added. The resulting solvated binary and ternary systems were used to generate topology and coordinates files which were used for carrying out molecular dynamics using Amber20 software [51]. Once the binary and ternary systems are prepared, the simulation workflow involves the following steps: minimization, equilibration and the production. The minimization step is carried out to remove any steric constraints and hot spots in the systems. The equilibration run is carried out in an isothermal isobaric ensemble (constant temperature and constant pressure simulation) to bring the system temperature to 300 K and 1 atmospheric pressure. The time scale for the equilibration run was kept at 20 ns. The production MD run again in an isothermal isobaric ensemble was carried out for 500 ns, using the final configuration from the equilibration simulation as the starting configuration. Post-simulation, trajectory analysis is performed using the CPPTRAJ module [52] to extract information about the system’s dynamics, including structural properties and energetics. In addition, the free energy calculations using molecular mechanics generalized Born surface area approach were carried out for the configurations corresponding to the last 50 ns from simulations.

## Results

### Microglia internalizes Tau in a time-dependent fashion

As mentioned early, microglia are primarily involved in constant surveillance of their microenvironment, phagocytosis of microbes, cellular debris, damaged cells and aggregated proteins [11]. In Alzheimer’s disease conditions, microglia phagocytose Tau species from the extracellular space by active endocytosis followed by degradation [22,21,53]. In our previous studies, we have demonstrated the internalization of recombinant Tau fragments by microglial cells in a time-dependent manner [22]. Here, we show microglial internalization of toxic Tau species, i.e. oligomers and aggregates in a time-dependent fashion **(Figure. 1A, Supplementary figure. 1).** N9 microglial cells were treated with Alexa-Fluor 647-labelled extracellular Tau species, monomer, oligomer and aggregates and analysed for internalization by immunofluorescence microscopic analysis. The internalization by microglial cells were quantified at different time intervals such as 5, 30, 60, and 180 minutes interval for monomer **(Figure. 1B),** oligomer **(Figure. 1C)** and aggregate **(Figure 1D).** These data clearly represents the rapid internalization of Tau monomers and oligomers as compared to aggregates. Following that, we confirmed the microglial internalization of Tau species by western blot analysis from lysed cells treated with extracellular Tau for increasing time-intervals. The level of internalized Tau species increased at different time intervals stained by Tau antibody, K9JA **(Figure 1E and 1F).** Tau monomer and oligomers shows clear internalization as compared to aggregates **(Figure. 1G)**.

**Figure 1.**
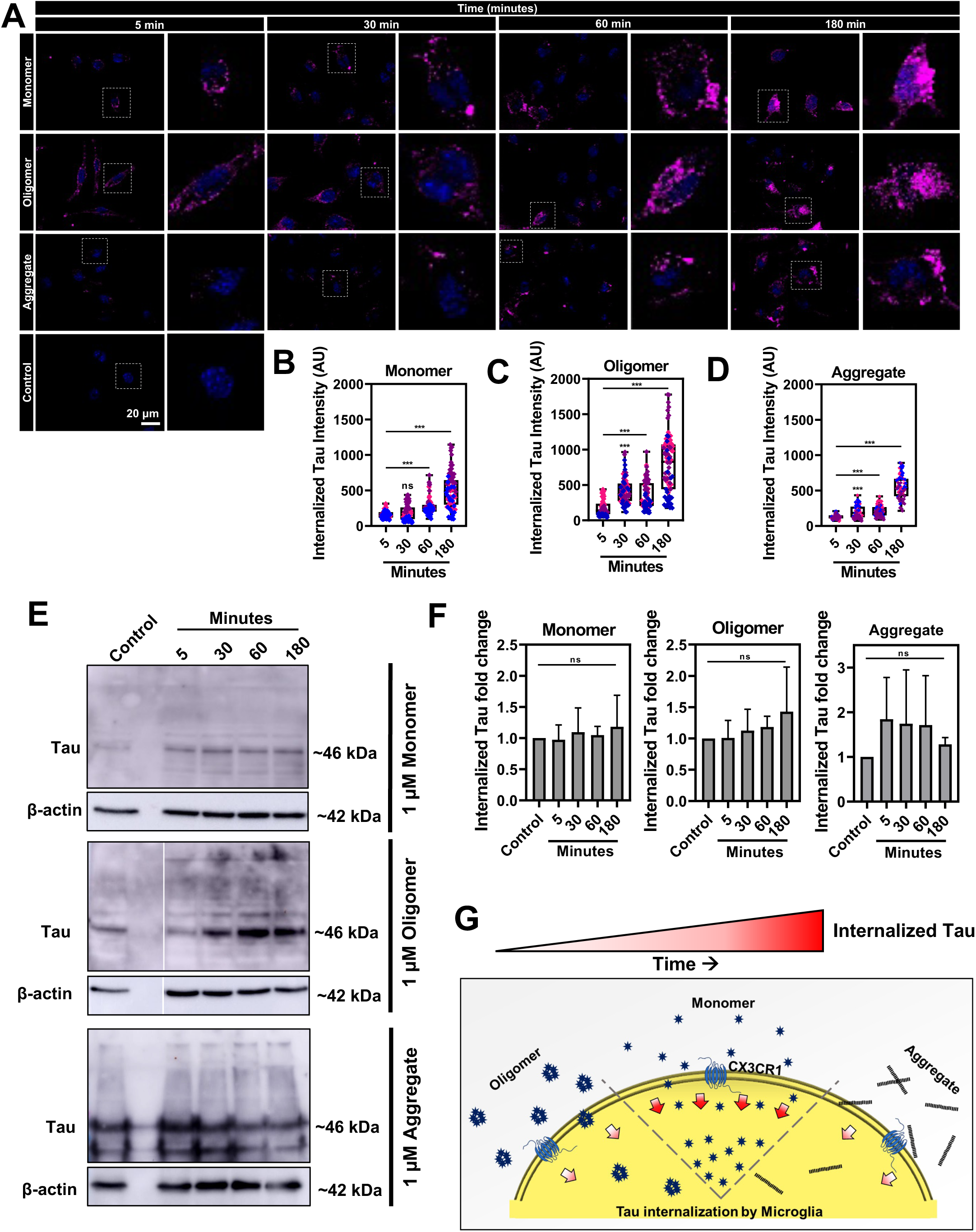
Time-dependent internalization of Tau species by microglia. A. Immunofluorescence microscopy analysis of Alexa Flour ^647^-labelled extracellular Tau internalization by N9 microglial cells at different time intervals 5, 30, 60 and 180 minutes. Extracellular Tau species were exposed at a concentration of 1 μM. Scale bar represents 20 µm. B, C, D. Quantification of internalized Tau intensity (n=75) of monomer (B), oligomer (C), and aggregate (D). Y-axis shows the internalized Tau intensity (arbitrary units) and the X-axis represents the different time interval observed for the study. E. Western blot analysis of Tau internalization by microglial cells. Extracellular Tau species (Monomer, Oligomer and aggregate) exposed cells (1 μM) were lysed and detected for Tau by western blot. Oligomer and aggregate blots show higher order proteins as well. β-actin is used as loading control for all blots. F. Quantification of western blot (n=3), normalized with β-actin loading control. G. Schematic representation of Tau internalization by microglia. One-way ANOVA is used for statistical analyses. The significance of multiple groups has been calculated by Tukey-Kramer’s post-hoc analysis for multiple comparisons at 5% level of significance and the p values are then calculated by comparing with control group. The level of significance is mentioned within the graph (*** p<0.001, ** p<0.01, * p<0.05, ns p≥0.05).

### Chemokine receptor CX3CR1 mediates Tau internalization in microglia

In 2017, *Bolos et al* illustrated the role of microglial CX3CR1 in Tau monomer internalization [12]. In this study, we are interested in exploring the role of this chemokine receptor, CX3CR1 in Tau species internalization under pathological conditions that include oligomers and aggregates. N9 microglial cells are exposed to extracellular Tau species in the presence of CX3CR1 antagonist, AZD8797 and the chemokine domain of fractalkine to determine the involvement of microglial CX3CR1 in oligomers and aggregates internalization. Internalized Tau intensity is observed at different time intervals in the presence of AZD8797, fractalkine and compared with control groups. The intensity of Tau monomer decreased significantly after 5 minutes in AZD8797-treated group as compared to control **(Figure. 2A and B)**. Whereas, Tau oligomers and aggregates shows enhanced Tau internalization at early time-points which decreases later at 3 hours **(Figure. 2A, C and D).** Similarly, CX3CR1 ligand, the chemokine domain of fractalkine reduced Tau monomer internalization at earlier time-points and enhanced internalization only at 3 hours. This proves the competitive binding of Tau monomers to CX3CR1 receptor in the presence of ligand, fractalkine. Whereas increased Tau internalization was observed at all time-intervals in oligomer and aggregate treated groups **(Figure. 2A-D).** These data suggest the CX3CR1 receptor-mediated Tau oligomer and aggregate internalization in microglia **(Figure. 2E).**

**Figure 2.**
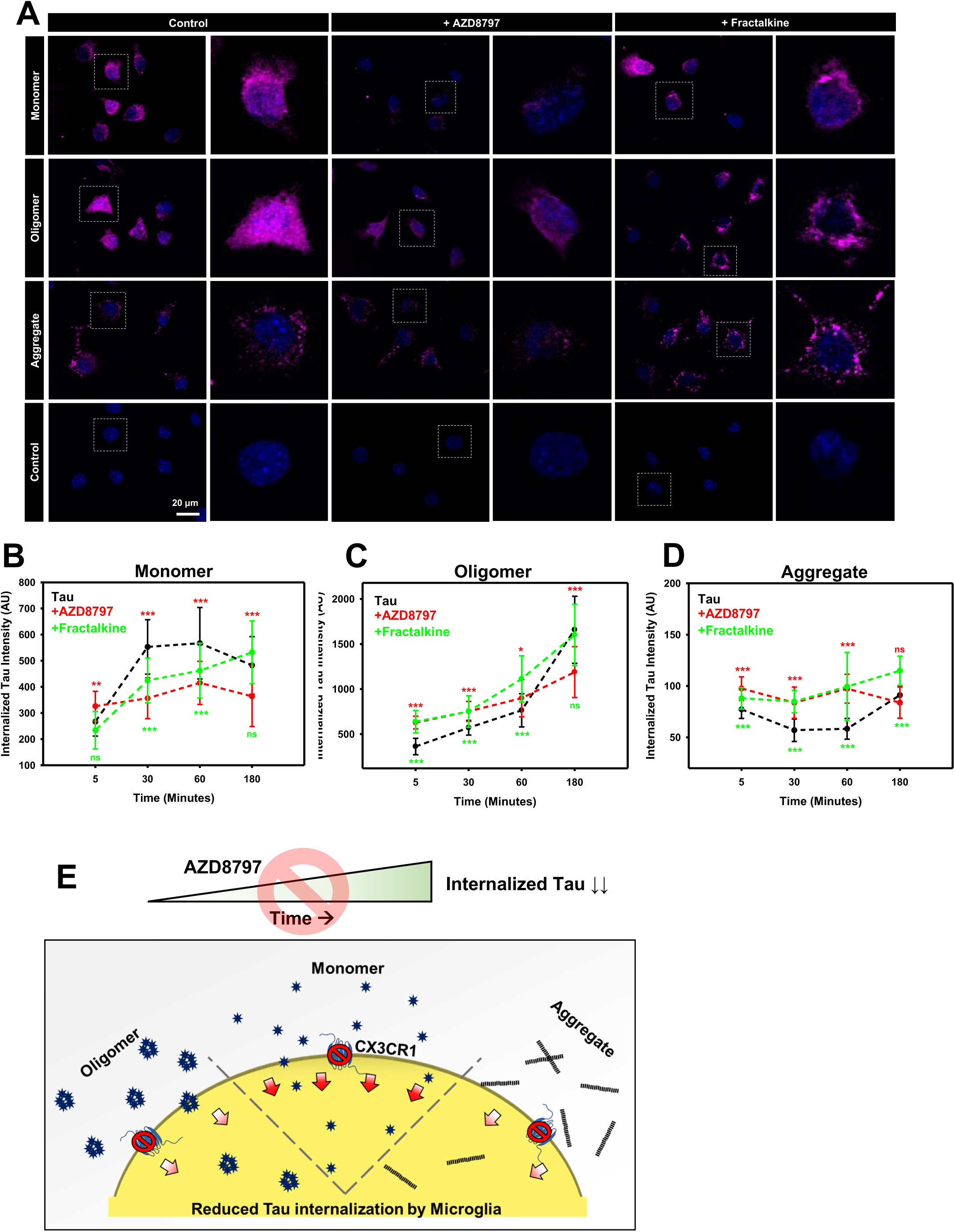
CX3CR1-mediated Tau internalization by microglia. A. Immunofluorescence microscopy analysis of Alexa Flour ^647^-labelled extracellular Tau internalization (1 μM) by microglial cells at 180 minutes in the presence of CX3CR1 antagonist and ligand, AZD8797 (100 nM) and fractalkine (chemokine domain) (250 ng/ml) respectively. Scale bar represents 20 µm. B, C, and D. Quantification of Tau monomer, oligomer and aggregate internalization respectively for the groups with AZD8797 and fractalkine treatment at 5, 30, 60 and 180 minutes (n=30). Y-axis shows the internalized Tau intensity (arbitrary units) and the X-axis represents the different time interval observed for the study. E. Schematic representation of CX3CR1-mediated Tau internalization. One-way ANOVA is used for statistical analyses. The significance of multiple groups has been calculated by Tukey-Kramer’s post-hoc analysis for multiple comparisons at 5% level of significance and the p values are then calculated by comparing with control group. The level of significance is mentioned within the graph (*** p<0.001, ** p<0.01, * p<0.05, ns p≥0.05). Red and green indicate the statistical significance of Tau internalization in AZD8797- and fractalkine-treated groups respectively in comparison with Tau control group of respective time points.

### Tau species act as chemoattractant for CX3CR1 receptor in microglia

Being a chemokine receptor, CX3CR1 bears an ability to sense varied range of ligands, including Tau and shows concentration-dependent migration towards it [12,27]. In this study, we are focused on understanding the dependency of CX3CR1 signalling during chemokine migration of microglia towards Tau protein. To comment on the CX3CR1 signalling, we treated microglia cells with Tau species and AZD8797 inhibitor, which blocks the CX3CR1 signalling for 24 hours. Out of all, Tau oligomer and aggregates found to be better chemoattractants as compared to Tau monomer, which showed significant wound healing after 24 hours. Similarly, inclusion of AZD8797 increased the rate of wound closure in the Tau oligomer and aggregates exposed condition rather than decreasing the migration of microglia **(Figure. 3A and B)**. The fact of AZD8797 as an allosteric non-competitive drug for CX3CR1 signalling, explain the possible increase of the rate of migration irrespective of inhibitor treatment [54]. Also, higher concentration of AZD8797 could be used to assess the capability of CX3CR1 to migrate towards Tau species in a chemotactic manner **(Figure. 3C)**.

**Figure 3.**
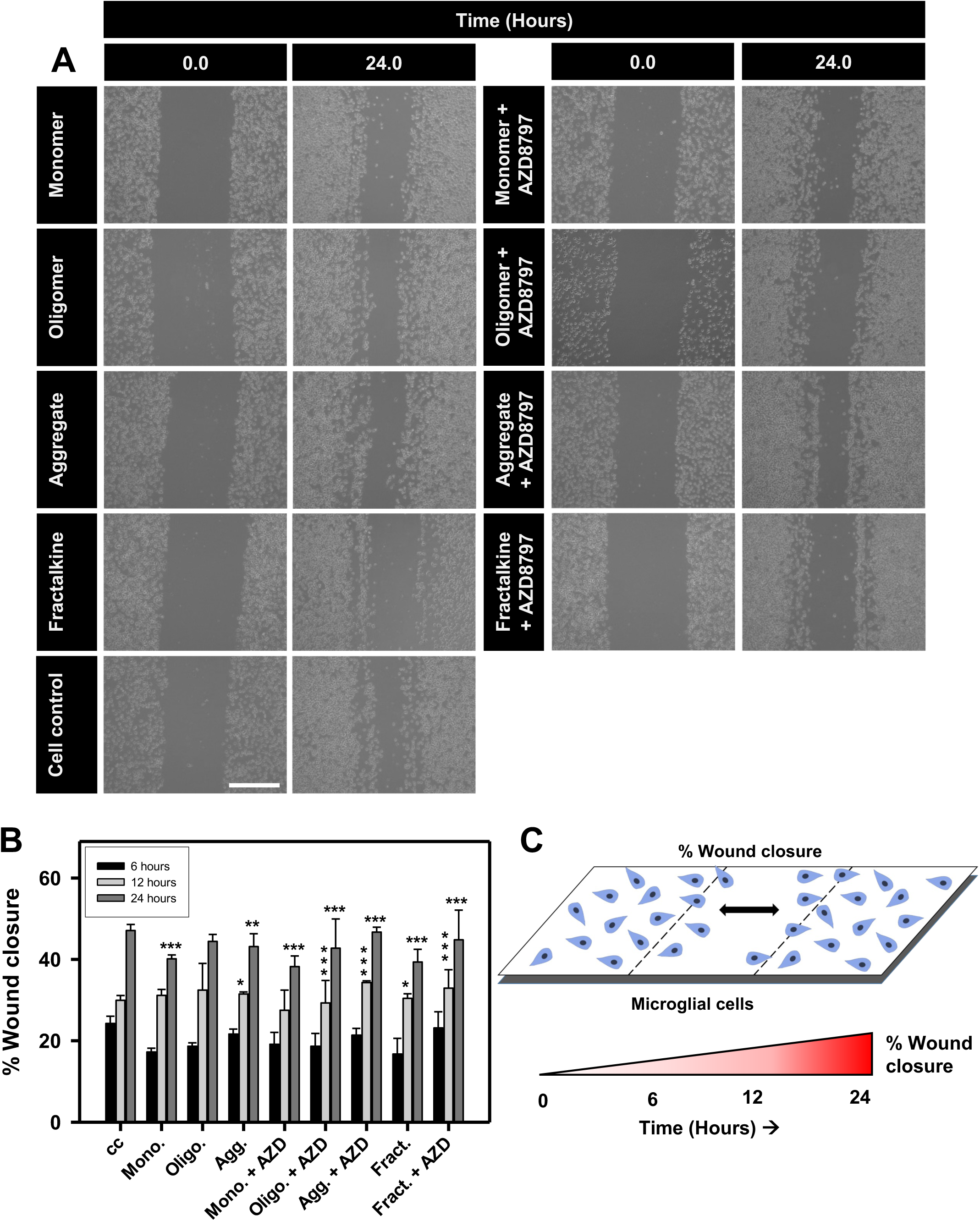
CX3CR1-dependent migration of microglia for extracellular Tau. A. Wound-scratch assay was performed to measure the migration capacity of microglia upon Tau and AZD8797 pharmacological inhibitor of CX3CR1. In N9 microglia cells, percentage wound-closure was evaluated at a regular interval of 6, 12, 24 hours and observed at 10X magnification through phase contrast microscope. Migration of cells into the wound was compared to wound created at 0 hour. Scale bar represents 100 μm. B. % wound closure was calculated at 6, 12, 24 hours and compared with cell control (no treatment) (n=30). C. Schematic diagram of microglial wound closure assay. One-way ANOVA is used for statistical analyses. The significance of multiple groups has been calculated by Tukey-Kramer’s post-hoc analysis for multiple comparisons at 5% level of significance and the p values are then calculated by comparing with control group (0 hour). The level of significance is mentioned within the graph (*** p<0.001, ** p<0.01, * p<0.05, ns p≥0.05).

### Microglial chemokine receptor, CX3CR1 interacts with extracellular Tau

As previously studied by Bolos *et al*, monomeric Tau competes with fractalkine for its binding to microglial chemokine receptor, CX3CR1 [12]. We were interested in studying the binding capability of Tau oligomers and aggregates to this chemokine receptor, CX3CR1. In this part of our study, N9 microglial cells were treated with extracellular Tau species, monomer, oligomer and aggregates and analysed for interaction and internalization with CX3CR1 receptor by immunofluorescence microscopic analysis. Alexa-flour 647 labelled Tau co-localizes with the microglial chemokine receptor, CX3CR1 at all the time-intervals denoting the direct receptor interaction. At early time points, Tau colocalization is observed at membrane region, whereas later time points showed cytosolic colocalization **(Figure. 4A, Supplementary figure. 2)**. Intensity profiling at the site of colocalization provides similar intensities for both channels, Tau and CX3CR1. The corresponding quantification of colocalization also supported the Tau-CX3CR1 interaction. Pearson’s coefficient of colocalization shows maximum colocalization at early time points for monomers and oligomers. Tau colocalization increases and reaches maximum at later time points in aggregate-treated group **(Figure. 4B)**. Further, we confirmed the interaction of Tau species by co-immunoprecipitation. Anti-Tau antibody-bound agarose beads were used to immunoprecipitate Tau species and their interacting partners. The samples are immunoblotted with CX3CR1 antibody followed by anti-Tau K9JA antibody. Co-immunoprecipitation clearly states the interaction of CX3CR1 receptor with Tau monomers, oligomers and aggregates **(Figure. 4C)**. The corresponding quantification shows maximum interaction with Tau monomer followed by aggregates and least interaction in case of Tau oligomers **(Figure. 4D)**. These data conclude the interaction of pathological Tau species, i.e. oligomers and aggregates with the microglial chemokine receptor, CX3CR1.

**Figure 4.**
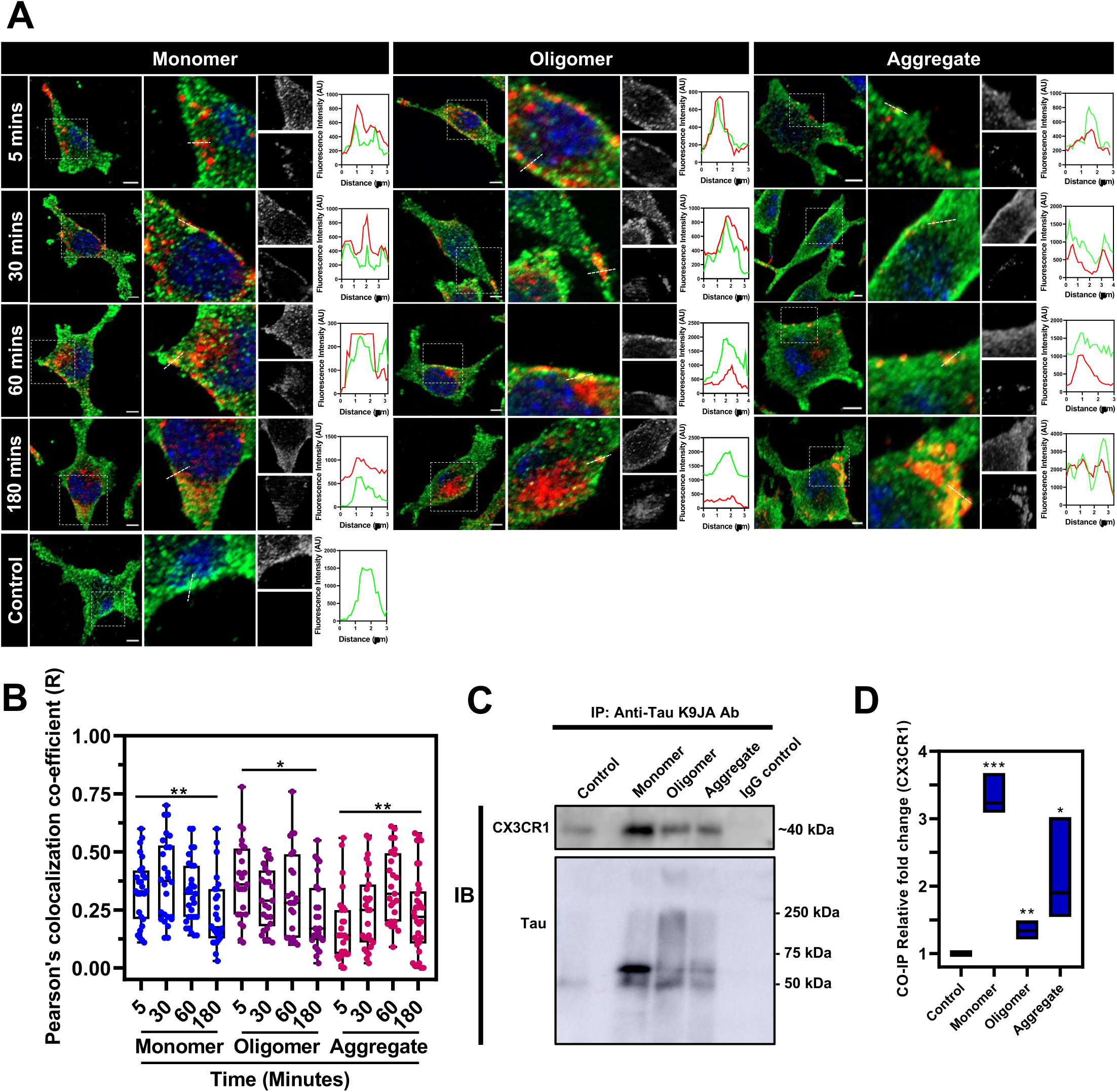
Tau species interaction with microglial CX3CR1 receptor. A. Immunofluorescence microscopy of N9 microglial cells showing colocalization between internalized Tau (Alexa Flour 647 labelled) (red) and CX3CR1 receptor (green) confirming Tau interaction with CX3CR1 receptor at different time intervals 5, 30, 60 and 180 minutes. Single cell representation, zoomed field of interest showing colocalization, grey scale images of green (top) and red (bottom) channels, and the corresponding intensity profile of the colocalizing field have been represented individually for every time-point of the study. The same has been studied in monomer, oligomer and aggregate individually. Scale bar represents 5 µm. The Y-axis of the intensity profile graph shows the fluorescence intensity (arbitrary units) and the X-axis shows the length (µm) of the profiling line (Profile line has been mentioned in the zoomed field of interest of corresponding images). B. The extent of colocalization is quantified using Pearson’s colocalization co-efficient (R) showing interaction at all the time intervals (n=30). One-way ANOVA is used for statistical analyses. The significance of multiple groups has been calculated by Tukey-Kramer’s post-hoc analysis for multiple comparisons at 5% level of significance and the p values are then calculated by comparing with control group. The level of significance is mentioned within the graph (*** p<0.001, ** p<0.01, * p<0.05, ns p≥0.05). **C.** CX3CR1 interaction with Tau species is confirmed by co-immunoprecipitation analysis. Tau species are immunoprecipitated with total Tau (K9JA) antibody and the interacting partners from the microglial cell lysate are co-eluted with Tau species. The samples are immunoblotted against CX3CR1 antibody and the same has been blotted with K9JA antibody. **D.** Quantification of CX3CR1 fold change (from CO-IP blot) compared between control and different Tau species (n=3). The statistical significance is determined by un-paired t-test, the p values are then calculated by comparing with control group and the level of significance is mentioned within the graph (*** p<0.001, ** p<0.01, * p<0.05, ns p≥0.05).

**Figure 5.**
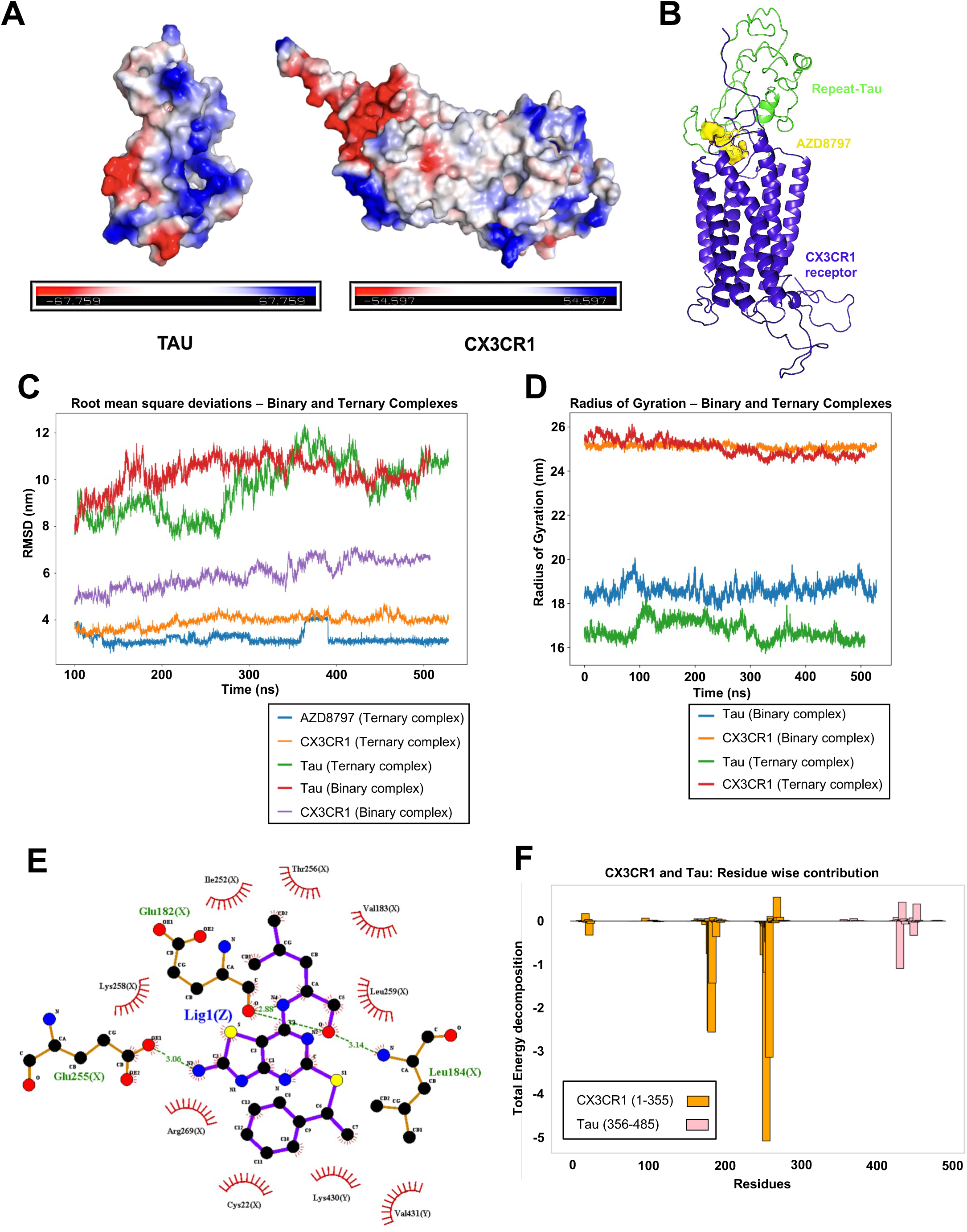
*In-silico* analysis of Tau-CX3CR1 interaction in the presence of CX3CR1 ligand, AZD8797 by Molecular docking and MD-simulation analysis. A. Representation of Electrostatic surface potentials of Tau and CX3CR1 receptor. Red region: Typically correspond to acidic residues or regions with an excess of electrons, suggesting negative electrostatic potential, Blue region: Generally correspond to basic residues or regions with a deficiency of electrons, suggesting positive electrostatic potential, White region: Correspond to regions with a balanced charge or no significant electrostatic potential. B. Ternary complex involving CX3CR1 protein, Tau protein and AZD8797 ligand. **C.** Graphical representation of the RMSD of CX3CR1, Tau, and the ligand (AZD8797) throughout the simulation time in different complexes. **D.** Graphical representation of the Radius of gyration of CXCR1 and Tau throughout the simulation time in different complexes. **E.** Ligplot diagram illustrating the interaction of CX3CR1, AZD8797 and Tau in Ternary Complex. **F.** Graphical illustration of residues that made a significant contribution to total energy decomposition for CX3CR1 and Tau. Residues from 1-355 is of CX3CR1 Receptor and 356-485 is Tau Protein.

### Computation analysis of Interaction of CX3XR1, Tau and AZD8789 ligand

In this study, we utilized Modeller to construct the structure of CX3CR1, selecting the model with the lowest DOPE score as it represents the most energetically favorable conformation. Next, we utilized CX3CR1 and Tau structure to calculate the electrostatic potential surface (ESP) and their charges were -1 and 10 respectively. The CX3CR1 has an overall negative electrostatic potential while Tau has an overall positive electrostatic potential suggesting the electrostatic complementarity between these two molecules that can be the reason behind the stable complex formation as shown in **Figure. 5A**. The regions of positive charge on Tau are interacting favorably with regions of negative charge on CX3CR1 and vice versa (shown in blue and red regions as observed in the contact area of CXCR1 and Tau protein). The complex formation was also evaluated from the docked structure containing CX3CR1, Tau and AZD8797 **(Figure. 5B)** using molecular dynamics simulation and free energy calculations. The convergence of various properties such as Root mean square deviation (RMSD) and Radius of gyration (Rg) for CX3CR1 and Tau were computed using trajectories which showed that both binary and ternary complex systems were stable during the entire course of simulation. **Figure 5C and D** show the evolution of these properties for both systems as a function of time. Our analysis revealed intriguing insights into the interaction between the molecules: the CX3CR1 receptor and the AZD8797 ligand exhibited the lowest RMSD values in the ternary complex, indicating a stable interaction between the ligand and the receptor in the presence of Tau. Tau protein displayed the highest RMSD values in both the binary and ternary complexes, indicating greater structural fluctuations which is due to its intrinsic nature of conformational flexibility **(Figure. 5C)**. Further, in-depth analysis of the radius of gyration (Rg) for over 5000 frames in both the binary and ternary complexes revealed that the CX3CR1 receptor exhibited the highest Rg values compared to Tau protein **(Figure. 5D)**. This suggests that the receptor adopts a more extended or elongated conformation, potentially due to its larger size. Interestingly, in the ternary complex, Tau protein displayed the lowest Rg values compared to the binary complex. This indicates that Tau adopts a more compact or folded conformation in the presence of the AZD8797 ligand and Tau may undergo structural rearrangements or exhibit modulated energetics when bound to the inhibitor bound receptor CX3CR1. Hence, it justifies the fact that presence of AZD8797 might alter the binding or signaling between Tau and CX3CR1. The conformational changes induced by AZD8797 in Tau may influence its interaction with CX3CR1, potentially affecting downstream signaling pathways or cellular processes associated with Tau pathology.

Further, the intermolecular hydrogen bonding analysis provides valuable insights into the structural dynamics of the binding interface between Tau and CX3CR1 receptors in both the binary and ternary complexes **(Figure. 5E, and Supplementary figure. 4).** In the binary complex, intermolecular hydrogen bonding involving residues GLU19, VAL182, and GLY383 from CX3CR1 and residues SER428, GLY435, and ASP266 from Tau respectively were observed, with average distances ranging from 2.64 to 2.86 Å and fractional occupancies ranging from 0.30 to 0.38. Hence, immunofluorescence assay, co-immunoprecipitation experiments as well as these intermolecular hydrogen bonding analysis **(Table. 1)** confirms the Tau protein interaction with CX3CR1. In ternary complex, hydrogen bonds between CX3CR1 and Tau occurred at average distances ranging from 2.62 to 2.87 Å, with fractional occupancies ranging from 0.32 to 0.44, while hydrogen bonds between the ligand and CX3CR1 involved residues GLU182, LEU184, and GLU255 from CX3CR1 with average distances ranging from 2.81 to 2.90 Å and fractional occupancies ranging from 0.28 to 0.52 **(Figure. 5E)**. Hence, these findings indicate that the ligand is forming stronger hydrogen bonds with the receptor (based on the increased hydrogen bond residence time). Moreover, in the ternary complex the lowered fractional occupancy of hydrogen bonds between Tau and CX3CR1 **(Table. 2)** comparative to the binary complex to ligand and CX3CR1 **(Table. 3)** suggests altered interaction between Tau and receptor due to the presence of ligand. The changes in the hydrogen bonding interaction pattern between the Tau receptor due to ligand underscores the ligand’s influence on Tau-CX3CR1 interactions, potentially impacting downstream signaling pathways relevant to Tau pathology.

The provided results of binding free energy calculations in **Table. 4** reveal important insights into the interactions between AZD8797, Tau, and CX3CR1. A more negative ΔGbinding value indicates a stronger affinity between molecules. The binding free energy for AZD897 with the Tau receptor complex is -32.5 kcal/mol and the analysis of residue wise contribution to binding free energies suggest the ligand interacts majorly with the receptor than that of Tau monomer **(Figure. 5F)**. The total residue-wise contribution value for CX3CR1 is -17.45 indicating a strong interaction with the ligands, suggesting a favorable binding affinity between CX3CR1 and the ligands AZD8797. This supports the observation that AZD8797 competes effectively with Tau for binding to CX3CR1, potentially inhibiting the internalization of pathological Tau species by microglial cells. The binding free energies of the Tau receptor complex in the presence of AZD8797 and absence shows the complexation energies are negative but the interactions are only modulated slightly due to the presence of the ligand. The receptor Tau binding free energies in presence and absence of ligand are - 47.4 kcal/mol and -50.9 kcal/mol respectively. This shows that the receptor Tau interactions are significant but altered significantly due to ligand which may affect the internalization of extracellular Tau.

### Tau species binding promotes CX3CR1 desensitization in microglia

Tau interaction with GPCR may alter the down-signalling cascades or promote receptor desensitization. Upon fractalkine binding, activated CX3CR1 receptor undergoes receptor desensitization mediated by GRK-2 phosphorylation and β-arrestin recruitment to the receptor [38]. β-arrestins are the scaffolding proteins that promote receptor desensitizing, internalization and activation of several other intracellular signals [55]. To prove the receptor desensitization and endocytosis, we studied the association of β-arrestin with the chemokine receptor, CX3CR1. N9 microglial cells treated with extracellular Tau species at different time-intervals such as 5, 30, 60 and 180 minutes are immunostained with CX3CR1 and β-arrestin. Time-dependent Tau treatment also shows CX3CR1 desensitization and colocalization with β-arrestin at initial time point of 5 minutes as compared to cell control which again reduces down during the later time periods **(Figure 6A)**. The quantification of colocalization by Pearson’s coefficient also supports the interaction of β-arrestin with CX3CR1 at 5 minutes which is maximum at monomer followed by oligomers and aggregates as compared to cell control **(Figure. 6B)**. This study supports the hypothesis of GRK-binding and receptor phosphorylation that ultimately leads to β-arrestin recruitment upon receptor activation by extracellular Tau exposure **(Figure. 6C)**.

**Figure 6.**
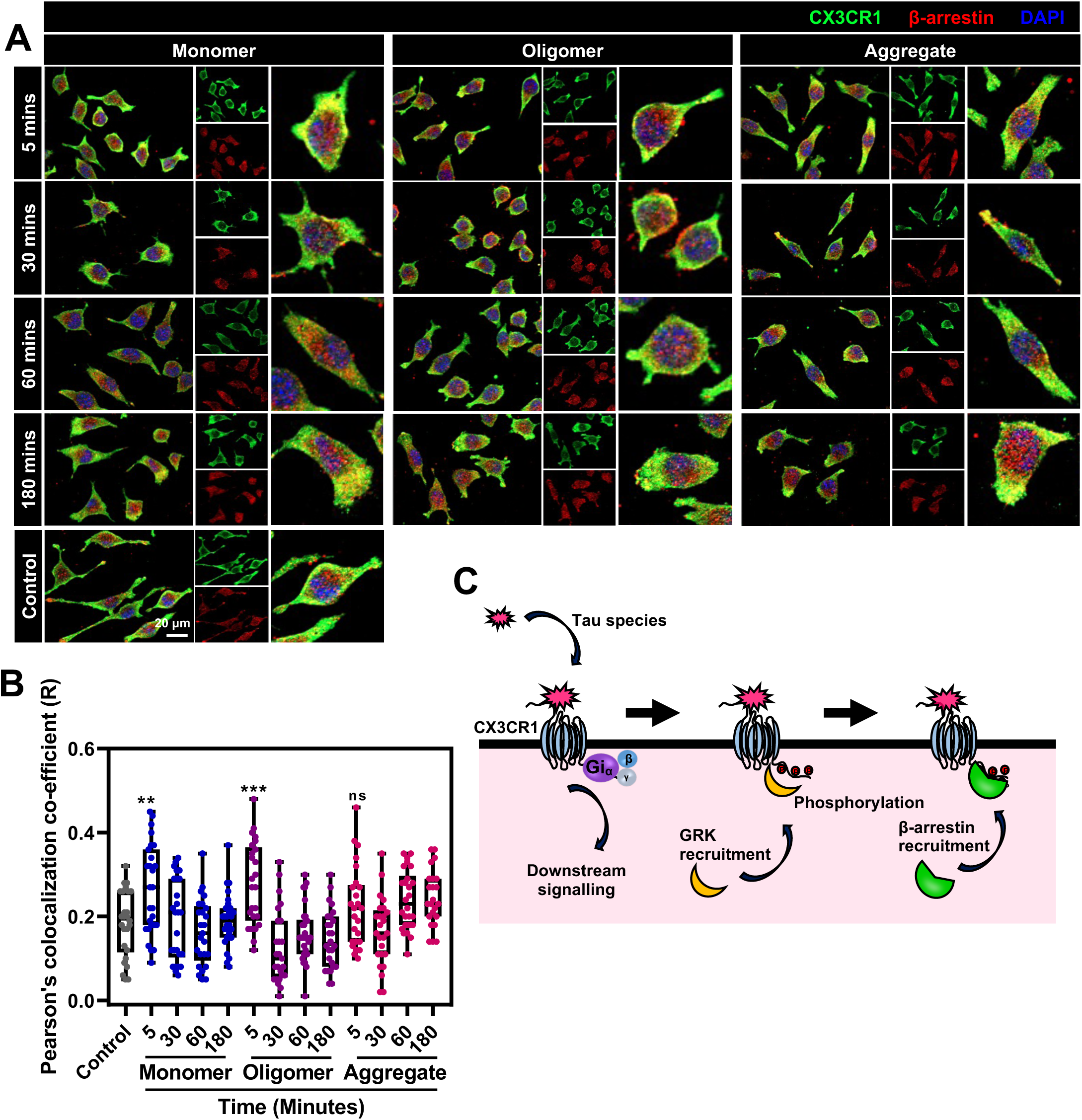
CX3CR1 desensitization upon Tau interaction in microglia. **A.** Immunofluorescence microscopy analysis of extracellular Tau (1 μM) treated N9 microglial cells at different time intervals 5, 30, 60 and 180 minutes. Cells were stained with CX3CR1 (green) and β-arrestin (red) showing colocalization in the treatment groups. Scale bar represents 20 µm. **B**. Corresponding quantification showing CX3CR1-βarrestin colocalization at initial 5 minutes as compared to cell control. Y-axis shows the Pearson’s coefficient of colocalization (R) and the X-axis shows Tau-treated groups at different time-intervals (n=50). One-way ANOVA is used for statistical analyses. The significance of multiple groups has been calculated at 5% level of significance and the p values are then calculated by comparing with control group. The level of significance is mentioned within the graph (*** p<0.001, ** p<0.01, * p<0.05, ns p≥0.05). **C**. Schematic representation of CX3CR1 receptor desensitization by receptor phosphorylation and β-arrestin recruitment.

### CX3CR1 receptor internalizes upon extracellular Tau exposure

G-protein coupled receptors upon ligand binding transmit signals from the cell surface. Following receptor activation, GPCRs internalizes to the endosomal compartments that further undergoes lysosomal degradation or recycles back to the membrane [56–58]. Hence, we studied the internalization pattern of CX3CR1 receptor and its distribution in N9 microglia upon extracellular Tau species at different time intervals such as 5, 30, 60 and 180 minutes from immunofluorescence assay **(Figure. 7A, Immunofluorescence images and Supplementary figure. 3)**. For this, we have performed the intensity profiling of individual cells at the cortical layer. Intensity profile of 6 µm was acquired from the membrane to cytosol of individual cells (n=50) and the surface to cytosolic graph was plotted for all Tau-treated groups **(Figure. 7A, Quantification graph)**. At cell control, the receptor intensity on cell surface remained unchanged. As compared to cell control, CX3CR1 intensity at membrane gradually decreases and shifts to cytosolic region with increase in time in all Tau-treated groups. In order to study the change in expression pattern of CX3CR1 receptor, we have performed western blot assay of total cell lysate of microglia upon extracellular Tau exposure at 24 hours **(Figure. 7B)**. The quantification graph normalized with β-actin expression shows insignificant changes in the total expression of CX3CR1 receptor upon Tau exposure **(Figure. 7C)**. This shows the clear change in the distribution of CX3CR1 receptor, i.e. the internalization of the receptor to endosomal compartments upon the binding of extracellular Tau species **(Figure. 7D)**.

**Figure 7.**
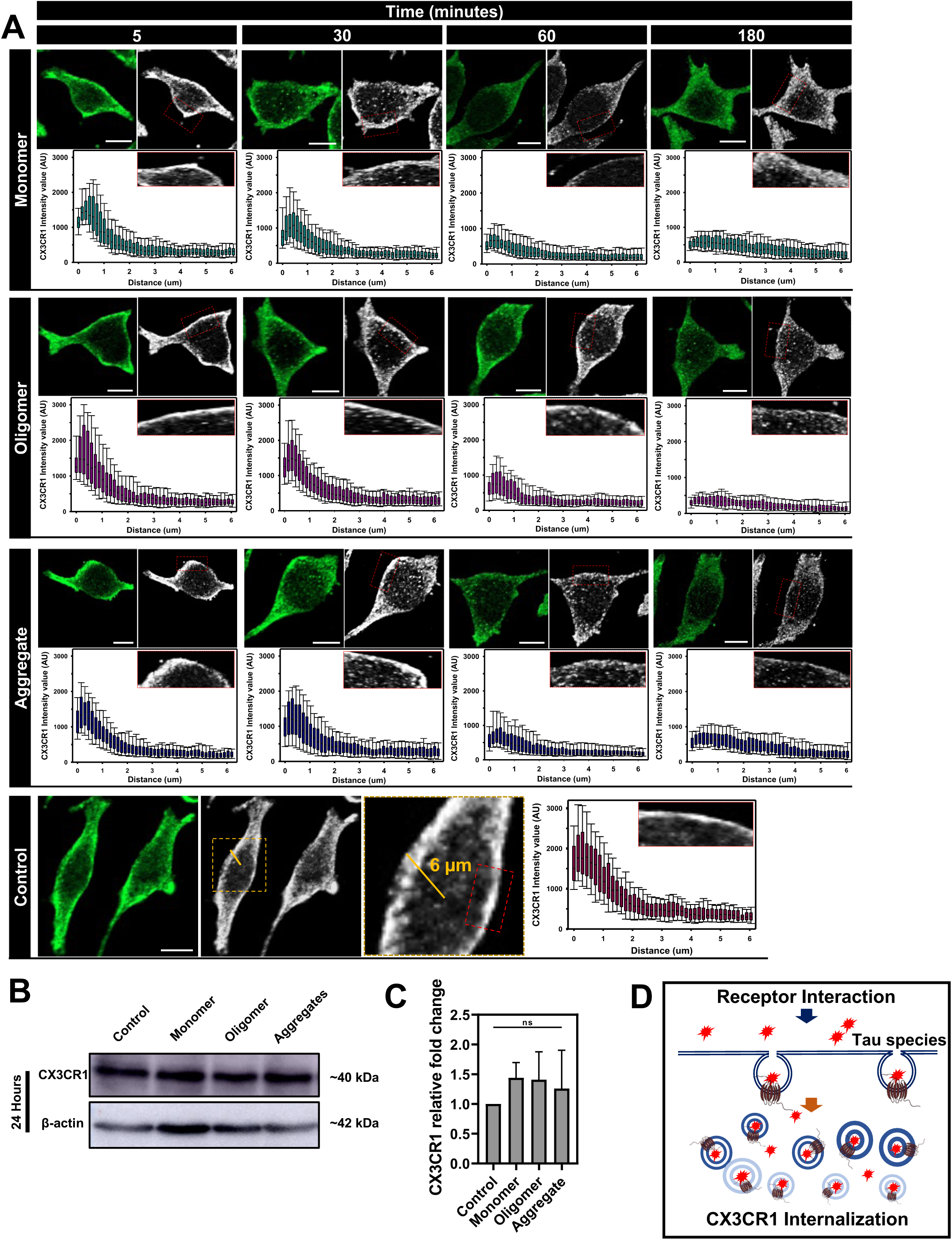
CX3CR1 distribution upon extracellular Tau exposure. **A.** Immunofluorescence microscopy analysis of extracellular Tau (1 μM) treated microglial cells at different time intervals 5, 30, 60 and 180 minutes. Cells were stained with CX3CR1 (green) in order to determine the localization of CX3CR1 receptor. The surface to cytosolic intensity ratio decreased upon increase in time is quantified by intensity profiling by Zen 2.3 software. CX3CR1 stained IF images (left) and grey scale pattern of CX3CR1 distribution (right) is provided on the top. The intensity profile of the CX3CR1 receptor in the cortical layer from membrane to cytosol is plotted for a distance of 6 µm. The Y-axis of the intensity profile graph shows the fluorescence intensity (arbitrary units) and the X-axis shows the length (µm) of the profiling line (yellow line shown in control image) (n=50). The intensity profile of individual time points have been represented at the bottom of IF images with the zoomed image showing the membrane to cytosol intensity of CX3CR1 receptor. Scale bar represents 5 µm **B.** Western blot analysis of CX3CR1 expression at 24 hours of extracellular Tau (1 μM) exposure. β-actin is used as loading control. **C.** The corresponding quantification of CX3CR1 expression (n=3), normalized with β-actin loading control. One-way ANOVA is used for statistical analyses. The significance of multiple groups has been calculated by Tukey-Kramer’s post-hoc analysis for multiple comparisons at 5% level of significance and the p values are then calculated by comparing with control group. The level of significance is mentioned within the graph (*** p<0.001, ** p<0.01, * p<0.05, ns p≥0.05). **D**. Diagrammatic representation of CX3CR1 interaction with extracellular Tau that leads to receptor endocytosis in microglial cells.

## Discussion

In this study, we have focussed on the microglial chemokine receptor, CX3CR1 and its interaction with different pathological Tau species, including oligomers and aggregates. We have provided evidences for Tau species interaction followed by receptor desensitization and internalization **(Figure. 8)**. In recent years, several receptors have been identified for interaction, signalling and internalization of Tau and Amyloid-β species in AD. Tau binding to GPCRs was discovered in early discovered in 2008 by Ramos *et al*. In neuronal cells, Tau directly binds to M1 and M3 muscarinic acetylcholine receptor and activates downstream signalling of these receptors [59,24]. The direct binding and activation is proved by transfecting M1 muscarinic acetylcholine receptor in COS-7 cells. Upon extracellular Tau exposure, the intracellular calcium level increased similarly compared to acetylcholine treated cells which is the native ligand of M1 muscarinic acetylcholine receptor. The direct binding of Tau to this receptor is further confirmed by immune-colocalization studies where Cy5 fluorescently labelled Tau completely colocalized with M1 muscarinic acetylcholine receptor in COS-7 cells. Further, both acetylcholine and Tau had a different binding sites and activation mechanisms. The acetylcholine binding desensitized the receptor whereas Tau binding could not induce desensitization in this receptor [59].

**Figure. 8.**
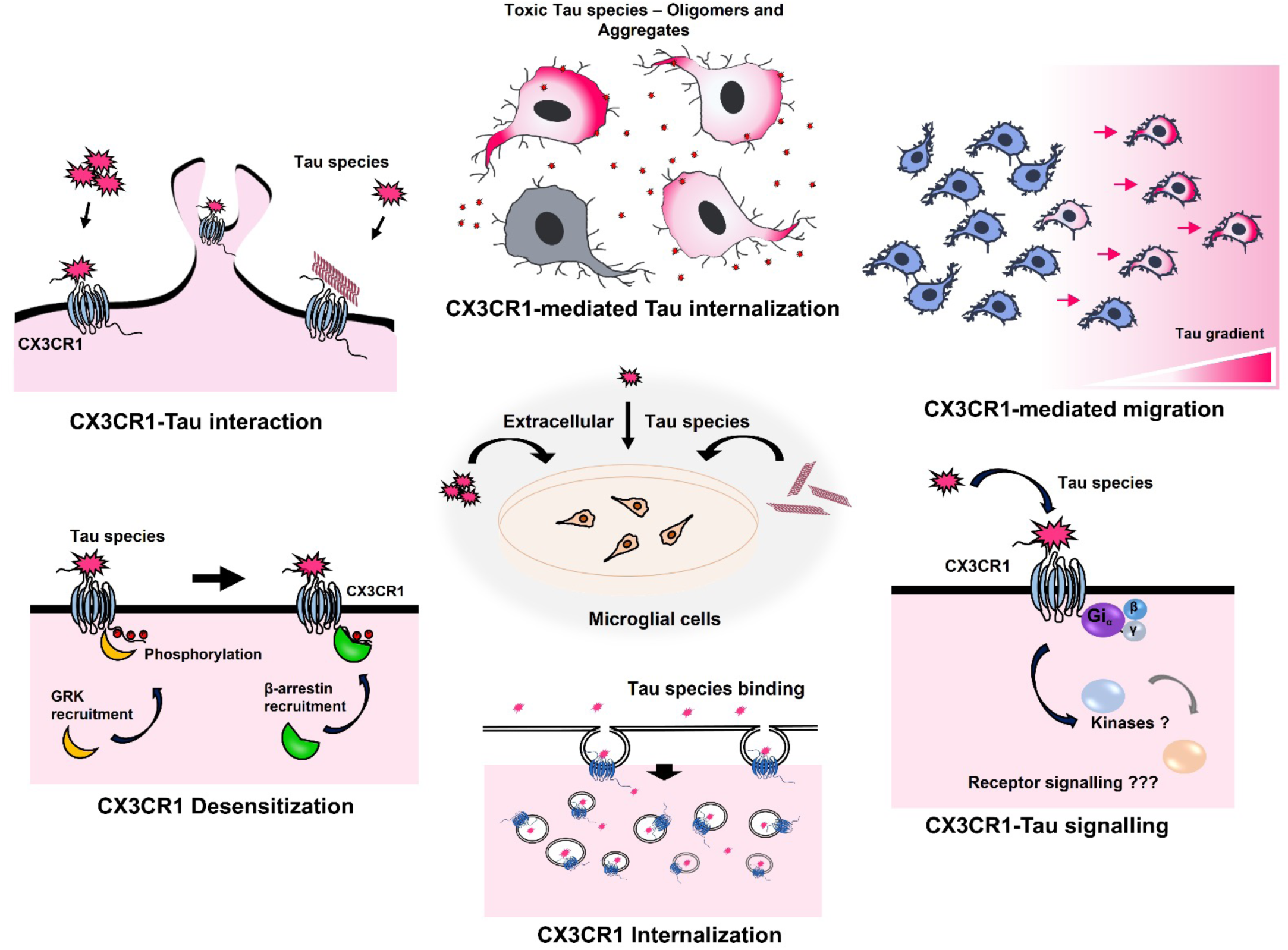
CX3CR1 – Role in extracellular Tau internalization. Extracellular exposure of Tau species promotes microglial activation and chemotactic migration. Tau species interacts with CX3CR1 receptor, desensitizes upon Tau binding, followed by β-arrestin recruitment and internalization of CX3CR1.

P2Y12R is a purinergic receptor abundantly expressed in microglia and aids in the surveillance for neuronal damage and neuroinflammation [60]. Microglia forms P2Y12R-mediated somatic synapses with neuronal cells to monitor the neuronal health [61]. Microglial P2Y12 receptor interacts with full-length Tau for its internalization and degradation [22,23]. Upon extracellular exposure of Tau species, microglia activated (increased Iba-1 expression) and showed P2Y12-mediated Tau internalization which is down regulated by pre-incubating with P2Y12 inhibitor, clopidogrel [22]. The repeat domain of Tau (Tau^RD^) is playing a key role in receptor binding which is further analysed by *in-silico* approaches such as molecular docking and MD-simulations [22]. The aggregated filaments also showed interaction which is further confirmed by co-immunoprecipitation and western blot from the total cell lysate of N9 microglial cells [23]. LRP1 is one such receptor responsible for Tau and Amyloid-β endocytosis followed by lysosomal trafficking in neuronal cells [62,25]. LRP1 receptor interacts with Tau, mediates Tau uptake and acts a key regulator of Tau spread and disease progression in Alzheimer’s disease. Genetic silencing of the LRP1 receptor significantly inhibited monomeric full-length Tau internalization in H4 neuroglioma cells [25]. LRP1 receptor knockdown also inhibited oligomeric species and to a lesser extent aggregated Tau fibrils internalization in neuronal cells [25].

Bolos *et al* reported that absence of microglial CX3CR1 receptor inhibits Tau internalization in microglia. Primary microglia cultures from wild-type and CX3CR1-knockout mice were exposed to 1 µM Tau and observed at different time intervals. CX3CR1 knockout microglia showed no internalization at early time intervals and a reduced internalization at 60 minutes as compared to wild-type groups. CX3CR1 knockout microglia also showed no change in the CD68+ expression pattern upon extracellular Tau exposure denoting the receptor role in microglial activation [12]. Monomeric Tau also acts as ligand for CX3CR1 and competes with fractalkine for its binding with the receptor [12]. In continuation of this study, we were interested in understanding the role of microglial CX3CR1 receptor under pathology conditions in the presence of Tau oligomers and aggregates. We have provided evidences for Tau species interacting with CX3CR1 receptor by immunofluorescence and co-immunoprecipitation. Migration of microglia stimulated by CX3CL1 is a part of homeostatic signalling in the brain environment. In AD condition, CX3CR1 mediated signalling and expression observed to be diminished in the presence of excessive pro-inflammatory cytokines IL-1β, TNF-α *etc.* [63,64]. Hence, understanding the extent of CX3CR1 activation by Tau species would be alternative approach to maintain the homeostatic signalling in disease condition. Tau species, particularly Tau oligomers and aggregates, possess a tendency to stimulate CX3CR1 signalling, widening a path to a potential treatment for AD. Our study clearly demonstrates the CX3CR1-mediated Tau internalization including oligomers and aggregates. As denoted by Bolos *et al*, fractalkine would have a competitive binding with CX3CR1 in the presence of Tau species that fluctuates its internalization as seen from the time-dependent graph in fractalkine-treated groups. From the wound-scratch assay, it is observed that Tau species, especially oligomers and aggregates promote migration similar to fractalkine. But, the inability of pharmacological inhibitor AZD8797 to reduce microglial migration in the presence of Tau, bears a possibility of stronger activation of CX3CR1 by Tau irrespective of the presence of antagonist. Moreover, our computational studies supported our immunofluorescence assay and co-immunoprecipitation experiments that confirms the Tau protein interaction with CX3CR1. *In-silico* studies suggest that AZD8797 competes effectively with Tau for binding to CX3CR1, potentially inhibiting the internalization of pathological Tau species by microglial cells. Despite the presence of AZD8797, the binding energy between Tau and CX3CR1 remains negative, indicating a favorable interaction. Additionally, in the ternary complex, the positive total residue-wise contribution to total binding free energies due to Tau is 0.11 kcal/mol which indicates that Tau do not have favorable interaction with the ligand in the ternary complex. As indicated in the experiments, AZD8789 strongly interacts with the receptor more than that of the Tau monomer. Therefore, the binding free energy results, along with the total residue-wise contribution values, collectively suggest that AZD8797, identified as a CX3CR1 antagonist, effectively binds to the receptor even in presence of Tau monomer modulating the internalization of pathological Tau species by microglial cells. The overall results suggest the propensity of extracellular Tau species as a ligand for CX3CR1 may lead to crosstalk in CX3CR1-fractakine axis that could ultimately lead to alternative downstream signalling pathways, enhanced microglial activation, *etc.* in AD conditions. Further research is required to understand the Tau-CX3CR1 downstream signalling pathways that enhances microglial activation, Tau species internalization and the endosomal trafficking of Tau-CX3CR1 upon internalization.

## Summary

In summary, we have studied the microglial chemokine receptor CX3CR1 mediated internalization of pathological Tau species, oligomers and aggregates from the extracellular space. This showed a time-dependent increase of internalized Tau and accumulation in microglial cells which is inhibited by CX3CR1 antagonist, AZD8797. We have proved the direct interaction of extracellular Tau species with the microglial chemokine receptor, CX3CR1 by immunofluorescence microscopy and co-immunoprecipitation analysis. We have further supported our studies by performing *in-silico* interaction analysis such as molecular docking, molecular dynamics and free energy calculations between Tau and CX3CR1 receptor. Further, the intensity profile analysis of immunofluorescence study showed the time-dependent receptor desensitization and endocytosis of CX3CR1 receptor following desensitization.

## Abbreviations

AD: Alzheimer’s disease
CNS: Central nervous system
GPCRs: G-protein coupled receptors
LRP: 1-Low density lipoprotein receptor-related protein 1
COS-7: CV-1 in Origin with SV40 genes-7
CD68: Cluster of Differentiation 68
IL-1β: Interleukin 1 beta
TNF-α: Tumor necrosis factor alpha. PDB – Protein Data Bank
MD: Molecular Dynamics
DOPE: Discrete Optimized Protein Energy
RMSD: Root mean square deviation
Rg: Radius of Gyration
ESP: Electrostatic surface potential.

## Acknowledgements

We are grateful to Chinnathambi’s lab members for their scientific discussions, helpful suggestions, and critical reading of the manuscript.

## Authors’ contributions

Hariharakrishnan Chidambaram and Subashchandrabose Chinnathambi performed the literature search and wrote the manuscript. Hariharakrishnan Chidambaram, Smita Eknath Desale and Subashchandrabose Chinnathambi performed the biochemistry and cell-biology experiments. Kapali Suri and Arulmurugan N performed the computational studies and wrote the manuscript. Subashchandrabose Chinnathambi conceived the idea of the work and supervised the project. All authors read and approved the final manuscript.

## Funding

The authors thank Department of Biotechnology for the fellowship. The internal infrastructure support from Department of Neurochemistry, National Institute of Mental Health and Neuro Sciences (NIMHANS), Bengaluru.

## Availability of data and materials

Not applicable.

## Declarations

### Consent for publication

The authors have no conflict of interest in the article.

### Competing interests

The authors declare that they have no competing interests.

## Notes

### Competing Interest Statement

The authors have declared no competing interest.

## References

1. Braak H, Braak E (1991) Neuropathological stageing of Alzheimer-related changes. Acta neuropathologica 82 (4):239–259

2. Braak H, Braak E (1991) Demonstration of amyloid deposits and neurofibrillary changes in whole brain sections. Brain pathology 1 (3):213–216

3. Hampel H, Schneider LS, Giacobini E, Kivipelto M, Sindi S, Dubois B, Broich K, Nistico R, Aisen PS, Lista S (2015) Advances in the therapy of Alzheimer’s disease: targeting amyloid beta and tau and perspectives for the future. Expert review of neurotherapeutics 15 (1):83–105

4. Binder LI, Frankfurter A, Rebhun LI (1985) The distribution of tau in the mammalian central nervous system. Journal of Cell Biology 101 (4):1371–1378

5. Sajjad R, Arif R, Shah A, Manzoor I, Mustafa G (2018) Pathogenesis of Alzheimer’s disease: role of amyloid-beta and hyperphosphorylated tau protein. Indian Journal of Pharmaceutical Sciences 80 (4):581–591

6. Neve RL, Harris P, Kosik KS, Kurnit DM, Donlon TA (1986) Identification of cDNA clones for the human microtubule-associated protein tau and chromosomal localization of the genes for tau and microtubule-associated protein 2. Molecular Brain Research 1 (3):271–280

7. Gong C-X, Liu F, Grundke-Iqbal I, Iqbal K (2005) Post-translational modifications of tau protein in Alzheimer’s disease. Journal of neural transmission 112 (6):813–838

8. Martin L, Latypova X, Terro F (2011) Post-translational modifications of tau protein: implications for Alzheimer’s disease. Neurochemistry international 58 (4):458–471

9. Sonawane SK, Chinnathambi S (2018) Prion-like propagation of post-translationally modified tau in Alzheimer’s disease: a hypothesis. Journal of Molecular Neuroscience 65 (4):480–490

10. Cowan M, Petri Jr WA (2018) Microglia: immune regulators of neurodevelopment. Frontiers in immunology 9:2576

11. Arcuri C, Mecca C, Bianchi R, Giambanco I, Donato R (2017) The pathophysiological role of microglia in dynamic surveillance, phagocytosis and structural remodeling of the developing CNS. Frontiers in Molecular Neuroscience 10:191

12. Bolós M, Llorens-Martín M, Perea JR, Jurado-Arjona J, Rábano A, Hernández F, Avila J (2017) Absence of CX3CR1 impairs the internalization of Tau by microglia. Molecular neurodegeneration 12 (1):59

13. Kleinberger G, Yamanishi Y, Suárez-Calvet M, Czirr E, Lohmann E, Cuyvers E, Struyfs H, Pettkus N, Wenninger-Weinzierl A, Mazaheri F (2014) TREM2 mutations implicated in neurodegeneration impair cell surface transport and phagocytosis. Science translational medicine 6 (243):243ra286-243ra286

14. Takahashi K, Rochford CD, Neumann H (2005) Clearance of apoptotic neurons without inflammation by microglial triggering receptor expressed on myeloid cells-2. The Journal of experimental medicine 201 (4):647–657

15. Perea JR, Bolós M, Avila J (2020) Microglia in Alzheimer’s disease in the context of tau pathology. Biomolecules 10 (10):1439

16. Leyns CE, Holtzman DM (2017) Glial contributions to neurodegeneration in tauopathies. Molecular neurodegeneration 12 (1):1–16

17. Spangenberg EE, Green KN (2017) Inflammation in Alzheimer’s disease: lessons learned from microglia-depletion models. Brain, behavior, and immunity 61:1–11

18. Evans LD, Wassmer T, Fraser G, Smith J, Perkinton M, Billinton A, Livesey FJ (2018) Extracellular monomeric and aggregated tau efficiently enter human neurons through overlapping but distinct pathways. Cell reports 22 (13):3612–3624

19. Kolay S, Vega AR, Dodd DA, Perez VA, Kashmer OM, White CL, Diamond MI (2022) The dual fates of exogenous tau seeds: Lysosomal clearance versus cytoplasmic amplification. Journal of Biological Chemistry 298 (6)

20. Perea JR, López E, Díez-Ballesteros JC, Ávila J, Hernández F, Bolós M (2019) Extracellular monomeric tau is internalized by astrocytes. Frontiers in neuroscience 13:442

21. Das R, Balmik AA, Chinnathambi S (2020) Phagocytosis of full-length Tau oligomers by Actin-remodeling of activated microglia. Journal of neuroinflammation 17 (1):1–15

22. Chidambaram H, Das R, Chinnathambi S (2022) G-Protein coupled Purinergic P2Y12 receptor interacts and internalizes TauRD-mediated by membrane-associated actin cytoskeleton remodelling in microglia. European Journal of Cell Biology:151201

23. Das R, Chinnathambi S (2021) Microglial remodeling of actin network by Tau oligomers, via G protein-coupled purinergic receptor, P2Y12R-driven chemotaxis. Traffic 22 (5):153–170

24. Gomez-Ramos A, Diaz-Hernandez M, Rubio A, Miras-Portugal M, Avila J (2008) Extracellular tau promotes intracellular calcium increase through M1 and M3 muscarinic receptors in neuronal cells. Molecular and Cellular Neuroscience 37 (4):673–681

25. Rauch JN, Luna G, Guzman E, Audouard M, Challis C, Sibih YE, Leshuk C, Hernandez I, Wegmann S, Hyman BT (2020) LRP1 is a master regulator of tau uptake and spread. Nature 580 (7803):381–385

26. Bjarnadóttir TK, Gloriam DE, Hellstrand SH, Kristiansson H, Fredriksson R, Schiöth HB (2006) Comprehensive repertoire and phylogenetic analysis of the G protein-coupled receptors in human and mouse. Genomics 88 (3):263–273

27. Chidambaram H, Das R, Chinnathambi S (2020) Interaction of Tau with the chemokine receptor, CX3CR1 and its effect on microglial activation, migration and proliferation. Cell & bioscience 10 (1):1–9

28. Meucci O, Fatatis A, Simen AA, Miller RJ (2000) Expression of CX3CR1 chemokine receptors on neurons and their role in neuronal survival. Proceedings of the National Academy of Sciences 97 (14):8075–8080

29. Imai T, Hieshima K, Haskell C, Baba M, Nagira M, Nishimura M, Kakizaki M, Takagi S, Nomiyama H, Schall TJ (1997) Identification and molecular characterization of fractalkine receptor CX3CR1, which mediates both leukocyte migration and adhesion. Cell 91 (4):521–530

30. Ostuni MA, Guellec J, Hermand P, Durand P, Combadière C, Pincet F, Deterre P (2014) CX3CL1, a chemokine finely tuned to adhesion: critical roles of the stalk glycosylation and the membrane domain. Biology open 3 (12):1173–1182

31. Harrison JK, Jiang Y, Chen S, Xia Y, Maciejewski D, McNamara RK, Streit WJ, Salafranca MN, Adhikari S, Thompson DA (1998) Role for neuronally derived fractalkine in mediating interactions between neurons and CX3CR1-expressing microglia. Proceedings of the National Academy of Sciences 95 (18):10896–10901

32. Hatori K, Nagai A, Heisel R, Ryu JK, Kim SU (2002) Fractalkine and fractalkine receptors in human neurons and glial cells. Journal of neuroscience research 69 (3):418–426

33. Garton K TACE (ADAM17) mediates the cleavage and shedding of Fractalkine (CX3CL1). J Biol Chem 10

34. Hundhausen C, Misztela D, Berkhout TA, Broadway N, Saftig P, Reiss K, Hartmann D, Fahrenholz F, Postina R, Matthews V (2003) The disintegrin-like metalloproteinase ADAM10 is involved in constitutive cleavage of CX3CL1 (fractalkine) and regulates CX3CL1-mediated cell-cell adhesion. Blood 102 (4):1186–1195

35. Hermand P, Pincet F, Carvalho S, Ansanay H, Trinquet E, Daoudi M, Combadière C, Deterre P (2008) Functional Adhesiveness of the CX3CL1 Chemokine Requires Its Aggregation ROLE OF THE TRANSMEMBRANE DOMAIN. Journal of Biological Chemistry 283 (44):30225–30234

36. Bhaskar K, Konerth M, Kokiko-Cochran ON, Cardona A, Ransohoff RM, Lamb BT (2010) Regulation of tau pathology by the microglial fractalkine receptor. Neuron 68 (1):19–31

37. Shafiei SS, Guerrero-Muñoz MJ, Castillo-Carranza DL (2017) Tau oligomers: cytotoxicity, propagation, and mitochondrial damage. Frontiers in aging neuroscience 9:83

38. Ge X-Y, Fang S-P, Zhou M, Luo J, Wei J, Wen X-P, Yan X-D, Zou Z (2016) TLR4-dependent internalization of CX3CR1 aggravates sepsis-induced immunoparalysis. American Journal of Translational Research 8 (12):5696

39. Chidambaram H, Chinnathambi S (2020) Role of cysteines in accelerating Tau filament formation. Journal of Biomolecular Structure and Dynamics:1–10

40. Šali A, Blundell TL (1993) Comparative protein modelling by satisfaction of spatial restraints. Journal of molecular biology 234 (3):779–815

41. Sonawane SK, Balmik AA, Boral D, Ramasamy S, Chinnathambi S (2019) Baicalein suppresses Repeat Tau fibrillization by sequestering oligomers. Archives of biochemistry and biophysics 675:108119

42. Tian W, Chen C, Lei X, Zhao J, Liang J (2018) CASTp 3.0: computed atlas of surface topography of proteins. Nucleic acids research 46 (W1):W363–W367

43. Honorato RV, Koukos PI, Jiménez-García B, Tsaregorodtsev A, Verlato M, Giachetti A, Rosato A, Bonvin AM (2021) Structural biology in the clouds: the WeNMR-EOSC ecosystem. Frontiers in molecular biosciences 8:729513

44. Van Zundert G, Rodrigues J, Trellet M, Schmitz C, Kastritis P, Karaca E, Melquiond A, van Dijk M, De Vries S, Bonvin A (2016) The HADDOCK2. 2 web server: user-friendly integrative modeling of biomolecular complexes. Journal of molecular biology 428 (4):720–725

45. Trott O, Olson AJ (2010) AutoDock Vina: improving the speed and accuracy of docking with a new scoring function, efficient optimization, and multithreading. Journal of computational chemistry 31 (2):455–461

46. Weiner PK, Kollman PA (1981) AMBER: Assisted model building with energy refinement. A general program for modeling molecules and their interactions. Journal of Computational Chemistry 2 (3):287–303

47. Wang J, Wolf RM, Caldwell JW, Kollman PA, Case DA (2004) Development and testing of a general amber force field. Journal of computational chemistry 25 (9):1157–1174

48. Frisch MJ, Trucks G, Schlegel H, Scuseria G, Robb M, Cheeseman J, Scalmani G, Barone V, Mennucci B, Petersson G (2009) Gaussian 09, rev. Gaussian Inc, Wallingford

49. Hornak V, Abel R, Okur A, Strockbine B, Roitberg A, Simmerling C (2006) Comparison of multiple Amber force fields and development of improved protein backbone parameters. Proteins: Structure, Function, and Bioinformatics 65 (3):712–725

50. Lee J, Cheng X, Jo S, MacKerell AD, Klauda JB, Im W (2016) CHARMM-GUI input generator for NAMD, GROMACS, AMBER, OpenMM, and CHARMM/OpenMM simulations using the CHARMM36 additive force field. Biophysical journal 110 (3):641a

51. Lee T-S, Allen BK, Giese TJ, Guo Z, Li P, Lin C, McGee Jr TD, Pearlman DA, Radak BK, Tao Y (2020) Alchemical binding free energy calculations in AMBER20: Advances and best practices for drug discovery. Journal of Chemical Information and Modeling 60 (11):5595–5623

52. Roe DR, Cheatham III TE (2013) PTRAJ and CPPTRAJ: software for processing and analysis of molecular dynamics trajectory data. Journal of chemical theory and computation 9 (7):3084–3095

53. Desale SE, Chinnathambi S (2021) α–Linolenic acid modulates phagocytosis and endosomal pathways of extracellular Tau in microglia. Cell adhesion & migration 15 (1):84–100

54. Cederblad L, Rosengren B, Ryberg E, Hermansson N-O (2016) AZD8797 is an allosteric non-competitive modulator of the human CX3CR1 receptor. Biochemical Journal 473 (5):641–649

55. Dorn GW (2009) GRK mythology: G-protein receptor kinases in cardiovascular disease. Journal of molecular medicine 87 (5):455–463

56. L Mohan M, T Vasudevan N, K Gupta M, E Martelli E, V Naga Prasad S (2012) G-protein coupled receptor resensitization-appreciating the balancing act of receptor function. Current molecular pharmacology 5 (3):350–361

57. Woolf PJ, Linderman JJ (2003) Untangling ligand induced activation and desensitization of G-protein–coupled receptors. Biophysical journal 84 (1):3–13

58. Calebiro D, Godbole A (2018) Internalization of G-protein-coupled receptors: Implication in receptor function, physiology and diseases. Best practice & research Clinical endocrinology & metabolism 32 (2):83–91

59. Gómez-Ramos A, Díaz-Hernández M, Rubio A, Díaz-Hernández JI, Miras-Portugal MT, Avila J (2009) Characteristics and consequences of muscarinic receptor activation by tau protein. European Neuropsychopharmacology 19 (10):708–717

60. Yu T, Zhang X, Shi H, Tian J, Sun L, Hu X, Cui W, Du D (2019) P2Y12 regulates microglia activation and excitatory synaptic transmission in spinal lamina II neurons during neuropathic pain in rodents. Cell death & disease 10 (3):1–16

61. Cserép C, Pósfai B, Lénárt N, Fekete R, László ZI, Lele Z, Orsolits B, Molnár G, Heindl S, Schwarcz AD (2020) Microglia monitor and protect neuronal function through specialized somatic purinergic junctions. Science 367 (6477):528–537

62. Fuentealba RA, Liu Q, Zhang J, Kanekiyo T, Hu X, Lee J-M, LaDu MJ, Bu G (2010) Low-density lipoprotein receptor-related protein 1 (LRP1) mediates neuronal Aβ42 uptake and lysosomal trafficking. PloS one 5 (7):e11884

63. Lauro C, Catalano M, Trettel F, Mainiero F, Ciotti MT, Eusebi F, Limatola C (2006) The chemokine CX3CL1 reduces migration and increases adhesion of neurons with mechanisms dependent on the β1 integrin subunit. The Journal of Immunology 177 (11):7599–7606

64. Zhou F, Liu R, Han P, Zhang X, Li Z, Zhang S, Liu C, Xia Y, Tang Z (2020) Pertussis toxin ameliorates microglial activation associated with ischemic stroke. Frontiers in Cellular Neuroscience 14:152

