## Supplementary material for "Microglia undergoes chemokine receptor, CX3CR1-mediated internalization of extracellular Tau": SI

### Supplementary figure 1

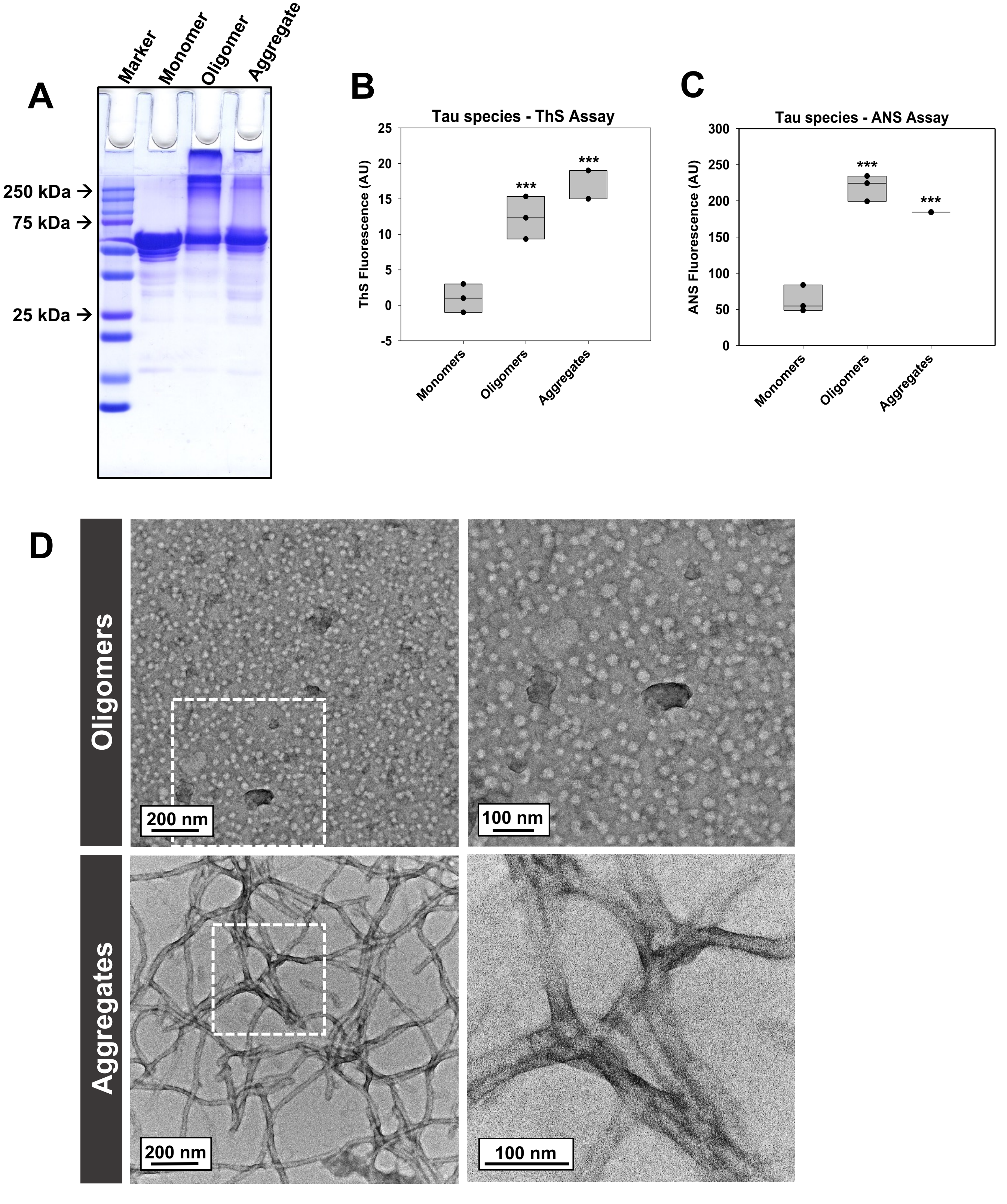


**Supplementary figure. 1. Characterization of Tau species. A.** 10% SDS-gel electrophoresis of Tau monomer, oligomer and aggregate.. **B** Thioflavin-S assay of Tau species- monomer, oligomer and aggregate. **C.** ANS assay of Tau species - monomer, oligomer and aggregate. **D.** Transmission electron microscopy images of Tau species - oligomer and aggregate.

### Supplementary figure 2

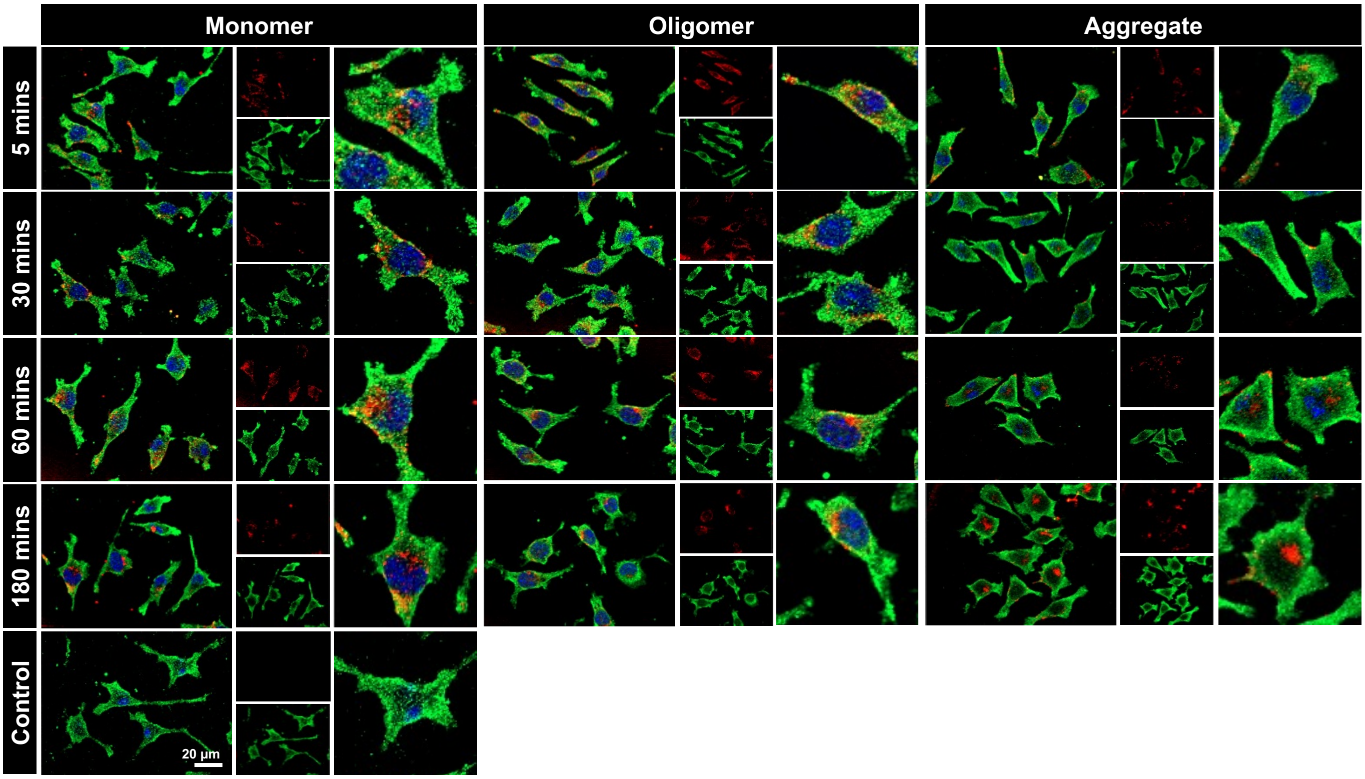


**Supplementary figure. 2. Tau species interaction with microglial CX3CR1 receptor.** Immunofluorescence microscopy of N9 microglial cells showing colocalization between extracellular Tau (Alexa Flour 647 labelled) (red) and CX3CR1 receptor (green) confirming Tau interaction with CX3CR1 receptor. Scale bar represents 20 µm.

### Supplementary figure 3

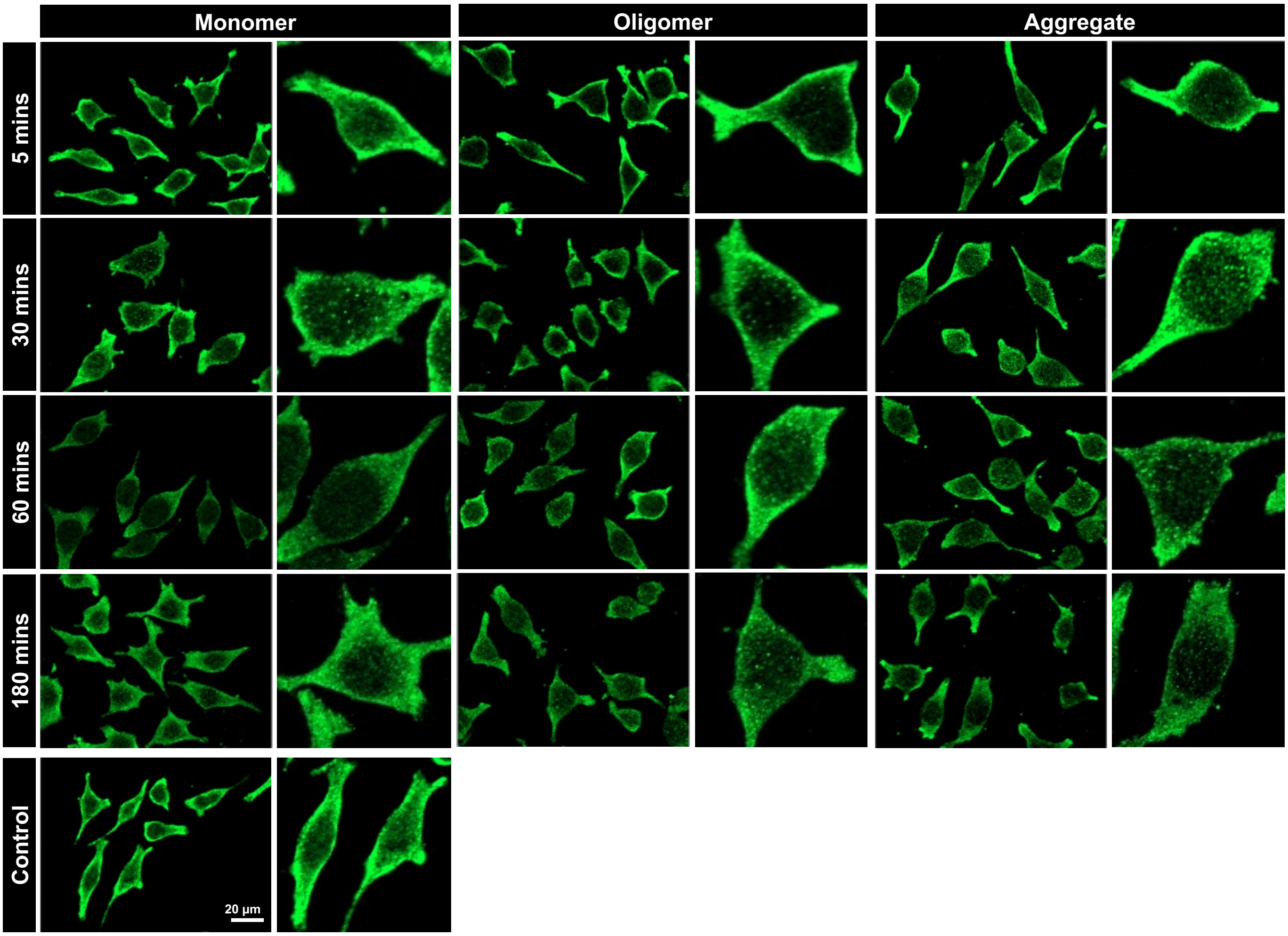


**Supplementary figure. 3. CX3CR1 distribution upon extracellular Tau exposure.** Immunofluorescence microscopy analysis of extracellular Tau treated microglial cells at different time intervals stained with CX3CR1 in order to determine the localization of CX3CR1 receptor. Scale bar represents 20 µm.

### Supplementary figure 4

A

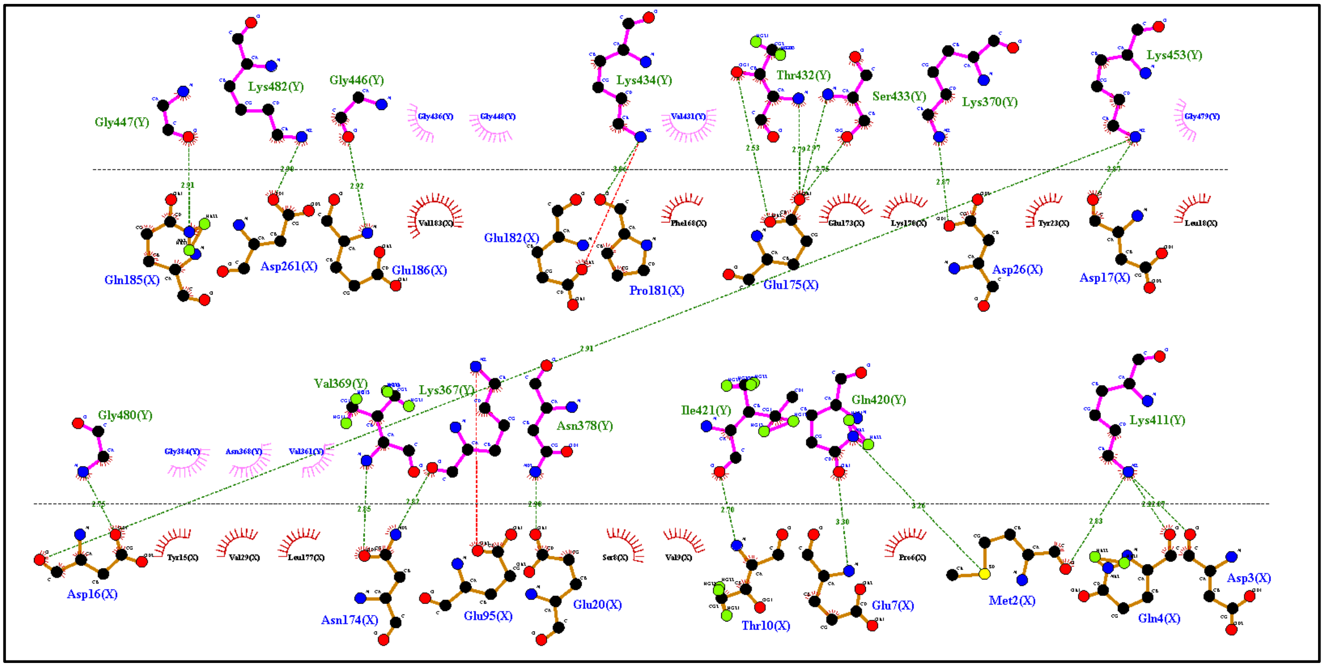


B

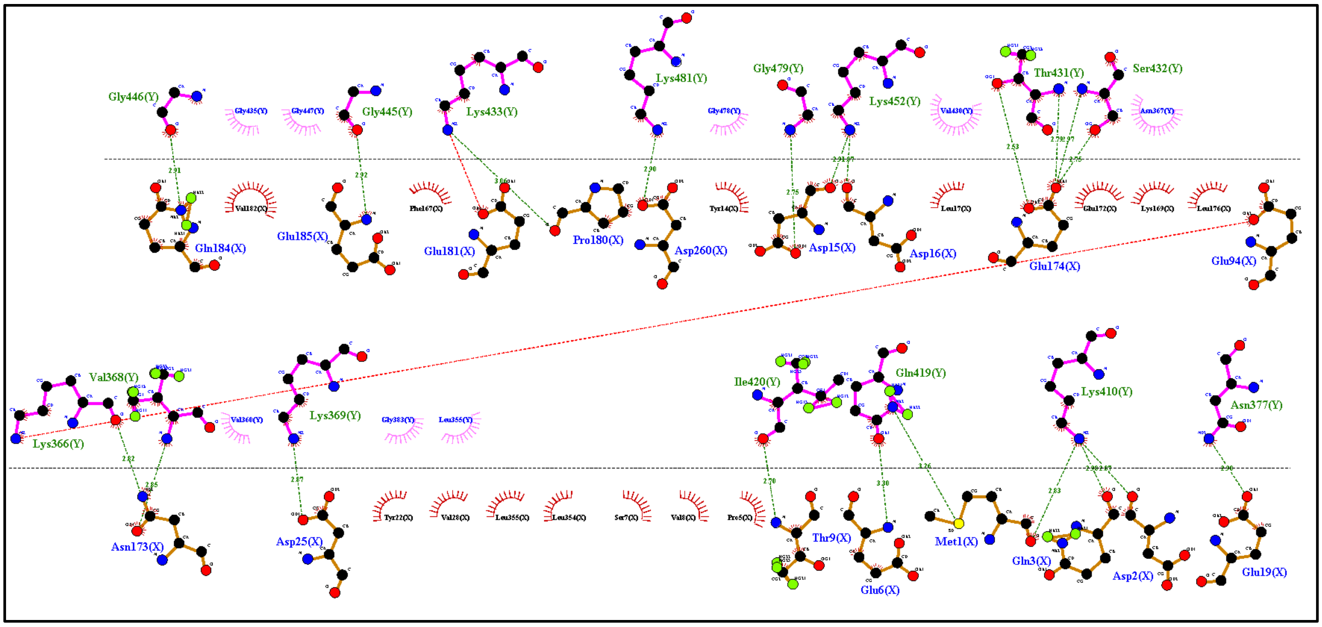


**Supplementary figure. 4. LigPlot interaction diagram. A.** Ligplot Diagram illustrating the interaction of CX3CR1 and Tau in Binary Complex. **B.** Ligplot Diagram illustrating the interaction of CX3CR1 and Tau in Ternary Complex.
